# A systematic review and meta-analysis of the prevalence of *Campylobacter* and non-typhoidal *Salmonella* along broiler chicken production and distribution networks

**DOI:** 10.64898/2026.11.15.766844

**Authors:** Chen Xin, Shanshan He, Nicola Di Girolamo, Casey L. Cazer, Priscilla F. Gerber

**Author notes:** Corresponding author: Priscilla F. Gerber, City University of Hong Kong, Tat Chee Avenue, Kowloon, Hong Kong.

## Abstract

*Campylobacter* spp. and non-typhoidal *Salmonella* (NTS) are leading bacterial causes of gastroenteritis worldwide and can be transmitted through contaminated broiler chicken products or contact with live birds and their environment. Asymptomatic carriage of these bacteria in broiler chickens enables pathogen shedding and subsequent product contamination at any component of broiler production and distribution networks (PDNs), including broiler farms, transport, slaughter, and retail. The study aimed to systematically review the published literature to 1) map the structure of broiler PDNs and identify where pathogens have been detected, 2) estimate the prevalence of *Campylobacter* spp. and NTS across broiler PDNs, and 3) assess the diversity and frequency of NTS serovars. A systematic search of the PubMed, Web of Science, and CABI Direct databases was conducted to identify peer-reviewed English-language articles reporting the prevalence of *Campylobacter* spp. or NTS across multiple components of broiler production and distribution. Separate multilevel meta-regression analyses were conducted for each pathogen to predict prevalence by PDN component and sample type, accounting for heterogeneity between studies and among estimates within studies. Frequency distributions were used to describe NTS serotype diversity. Data were pooled from 24 *Campylobacter* spp. studies and 23 NTS studies across 15 and 12 countries, respectively. Predicted *Campylobacter* spp. and NTS prevalence varied across PDN components and sample types, with the highest predictions at slaughter facilities. Predicted *Campylobacter* spp. prevalence reached 58.9% (95% CI, 29.0–83.4%) in pre-chill samples (after evisceration but before chilling), and NTS was 34.4% (95% CI, 17.6–56.2%) in broiler samples (intestinal contents, cecal and cloacal samples), which was significantly higher that on farms at the end of rearing (17.1%; 95% CI, 8.8–30.5%; OR = 2.54; *p* = 0.037). Infantis, Enteritidis, Kentucky, Hadar, and Typhimurium were the top five NTS serovars across all included studies. Predominant serovars were Enteritidis in farm broiler samples, Hadar in farm and slaughter environmental samples, and Infantis in retail product samples. These findings highlight slaughter facilities as hotspots with high *Campylobacter* spp. and NTS prevalence, supporting their prioritization for targeted monitoring and implementation of contamination-control measures. Given the variation in NTS serovar distribution, surveillance should include multiple sample types across PDN components.

## 1. Introduction

*Campylobacter* spp. and non-typhoidal *Salmonella* (NTS) are major bacterial causes of foodborne diarrheal disease worldwide (Havelaar et al., 2015). Poultry, including broiler chickens, can carry these pathogens without clinical signs (Newell & Fearnley, 2003; Shaji et al., 2023), which can contaminate the surrounding environment, carcasses, and raw chicken meat, creating opportunities for foodborne pathogen exposure (Alali & Hofacre, 2016). Human infection with *Campylobacter* spp. and NTS is strongly linked to handling, preparing, and consuming contaminated broiler chicken products, as well as contact with live birds and their environment (Davydova et al., 2025; EFSA Panel on Biological Hazards (BIOHAZ), 2011; Eisenstein et al., 2025; Hale et al., 2012). *Campylobacter jejuni* (*C. jejuni*) and *Campylobacter coli* (*C. coli*) are the main species responsible for human campylobacteriosis (Cody et al., 2017). Similarly, NTS, which encompasses all *Salmonella enterica* serovars except *S*. Typhi and *S*. Paratyphi (Fleckenstein et al., 2021), comprises more than 2,500 serovars (Grimont & Weill, 2007). Among these, *S*. Enteritidis was the most frequently reported serovar associated with human salmonellosis in the European Union and the United States, and was commonly detected in broiler flocks, chicken meat, and chicken carcasses (European Food Safety Authority and European Centre for Disease Prevention and Control (EFSA and ECDC), 2018; Walter et al., 2025).

Broiler chicken production and distribution networks (PDNs) connect farms, traders, transport operations, slaughter and processing facilities, live bird markets, and retail outlets, and these networks may be vertically integrated through company ownership or contractual coordination, or non-integrated, with independently operated components linked through trade (Hennessey et al., 2021; Thi Dien et al., 2023). Within PDNs, pathogens carried by live broilers on farms may contribute to downstream contamination of carcasses and raw chicken meat during handling and processing (Golden et al., 2021; Slader et al., 2002). Contaminated transport crates, slaughter equipment, and processing environments can provide additional sources of contamination (Kudirkienė et al., 2011; Rasschaert et al., 2006; Rasschaert, Houf, & De Zutter, 2007). These complex networks create many opportunities for broiler chickens, their products, and the surrounding environment to be contaminated with these foodborne pathogens. Therefore, mapping network components and knowing their connections provides a framework for investigating pathogen distribution and exploring potential routes of foodborne pathogen transmission. Sampling live broiler chickens, raw chicken products, and environmental samples across these components can reveal variation in the prevalence of *Campylobacter* spp. and NTS and help characterize differences in pathogen prevalence across PDN components. For example, a study in northern Vietnam estimated a higher probability of site-level NTS contamination at markets and slaughter facilities than on farms, based on environmental sampling (Xin et al., 2025). However, individual supply-chain studies differ in the components they cover and the sample types they examine, such as samples from broiler chickens, carcasses, raw chicken meat, or the environment. Synthesizing evidence across these studies, while distinguishing sample types, can therefore provide a broader picture of pathogen distribution across PDNs.

Previous systematic reviews and meta-analyses have characterized pre-harvest sources of *Campylobacter* spp. and *Salmonella* in poultry production (Wang et al., 2023a, 2023b). In the United States, Golden and Mishra et al. have also synthesized prevalence estimates across farming, processing, and retail samples, comparing conventional and alternative production systems (Golden & Mishra, 2020). However, to our knowledge, reviews providing these pathogen-specific prevalence estimates across PDN components and across sample types within each component remain limited.

Therefore, this systematic review and meta-analysis targeted scientific literature investigating *Campylobacter* spp. or NTS prevalence across multiple components of broiler PDNs and aimed to 1) map the structure of broiler PDNs from the end of on-farm rearing to retail and identify where pathogens have been detected based on included studies, 2) estimate pathogen-specific prevalence across PDN components and sample types using multilevel meta-regression analysis, and 3) characterize the distribution pattern of NTS serovars reported in the included studies.

## 2. Materials and methods

### 2.1. Study design

This systematic review was conducted in accordance with the PRISMA guidelines (Page et al., 2021). The review question addressed the prevalence of *Campylobacter* spp. and NTS in samples collected across components of broiler PDNs, from the end of the on-farm broiler rearing period to retail, and followed the population and outcome (PO) format (Cooke et al., 2012). The population of eligible sample types comprised biological samples from broiler chickens, chicken products, and associated environmental samples. The primary outcome was sample-level prevalence for the pathogen. A secondary outcome was the distribution of NTS serotypes among serotyped isolates.

### 2.2. Protocol and registration

The protocol was registered on the Open Science Framework (OSF) platform on 18 December 2025 (doi: 10.17605/OSF.IO/Y9AVQ). The protocol document was subsequently revised and finalized on 15 November 2026 and is available on the associated OSF project (https://doi.org/10.17605/OSF.IO/2N7JX). It detailed the research question, definition, eligibility criteria, timeframe, the subject areas of the studies, information source, search strategy, study records, quality assessment, and plan of data synthesis.

### 2.3. Literature search strategy

A systematic search was conducted in PubMed, Web of Science, and CABI Direct electronic databases in January 2026. All journal articles published in English were eligible. Boolean operators ‘OR’ and ‘AND’ were used to generate appropriate search strings. The PubMed database was queried using only the title and abstract. CABI databases were queried using only the abstract. The Web of Science was queried using the “Topic” search field (TS) to search within the title, abstract, and author keywords. The search strategies in PubMed, using keywords and search strings, are listed in Supplementary Table S1. The search strategies in CABI are shown in Supplementary Table S2; those in Web of Science are in Supplementary Table S3.

### 2.4. Eligibility and exclusion criteria

The included studies were peer-reviewed, English-language primary research articles that met all inclusion criteria detailed in Table 1. Notably, only supply-chain studies that reported pathogen prevalence across at least two PDN components, from the end of the on-farm broiler rearing period to the retail sale of raw broiler products, were eligible. No geographic restrictions were applied. Studies were excluded if they: 1) did not focus on *Campylobacter* spp. or NTS; 2) involved typhoidal *Salmonella* (Typhi or Paratyphi A, B, C); 3) restricted the detection to a single NTS serotype; 4) did not involve supply chain pathogen monitoring or were limited to a single component; 5) did not report overall sample prevalence, such as studies focusing solely on bacterial isolates or those conducting targeted sampling of known pathogen-positive broiler flocks; or 6) were experimental studies, case reports, or case series.

**Table 1.** Inclusion criteria that were applied to the papers retrieved from databases.

| **Inclusion criteria** | **Descriptions of the criteria** |
|---|---|
| Target animal | Broiler chickens |
| Target foodborne pathogens | *Campylobacter* spp., or NTS |
| Target samples<br>(biological samples) | Broiler chickens (cecal, cloacal samples, or other broiler samples); broiler chicken products (carcass, skin, feet, feather, meat, or other product samples) |
| Target samples (environmental samples) | Feces or litter, water, soil, or other environmental samples |
| Detection method | The study should mention the detection method of pathogen. |
| Outcome of interests | The study reports sample-level prevalence for the pathogen, including the original data of total sample size and number of positive samples. |

### 2.5. Data collection and extraction

The screening process was conducted from February 2026 to March 2026. Records extracted from each database were exported in PubMed or RIS format. Subsequently, these publications from three databases were imported to Covidence, a web-based platform designed for systematic review management (Veritas Health Innovation, 2023). Any duplicates across databases were identified and removed within Covidence.

Title and abstract screening were conducted independently by two reviewers (CX and SH), with CX serving as the lead reviewer. Articles that clearly failed to meet the inclusion criteria or satisfied any exclusion criteria were excluded at this stage. To enhance screening efficiency, the ‘Most relevant’ machine-learning feature in Covidence was utilized (Veritas Health Innovation, 2023), allowing the system to prioritize and display the studies most likely to be included first.

Subsequently, full-text copies were obtained for all potentially eligible studies and for those with unclear eligibility based on title and abstract review. For articles that were not open access or otherwise inaccessible, assistance was requested from Cornell University library staff. In the following reviewing phase, the full-text articles were independently screened by two reviewers (CX and SH). Disagreements were resolved through discussion. All reasons for exclusion at the full-text stage were recorded.

Once all eligible articles were identified, data were independently extracted from each paper by two reviewers (CX and SH), with discrepancies resolved through discussion and a final consensus reached. For each study, the following information was extracted: (1) publication details (title, authors, and publication year), (2) sampling-component characteristics (country, name of component, operational scale of the sampled component [i.e., small-scale or large-scale], year of the survey, source of samples, and laboratory method), (3) outcome (the total sample size and number of positive samples), and (4) NTS serotypes (with the number of isolates of each serotype) (Supplementary Table S4). The extracted information was entered into a Microsoft Excel spreadsheet for quality assessment and data preparation for analysis. For this review, components of the broiler PDN were classified according to supply-chain stage and setting in which samples were collected: (1) farms, including free-range, small-scale, and commercial broiler farms, where sampling was conducted near the end of the rearing period; (2) transportation, comprising transport vehicles and cages; (3) live-bird markets, comprising venues that sold live birds, including those in which vendors supplied traders for onward resale; (4) slaughter facilities, comprising sites where live broilers were processed into products, including small-scale slaughter points, large-scale slaughterhouses, and processing plants; and (5) retail, comprising sites of sale for raw broiler products, including retail markets, stalls, and supermarkets. Information extracted from eligible studies was synthesized to characterize component settings and their potential connection, thereby defining the structure of broiler PDNs. They were further mapped and classified as integrated or non-integrated based on the nature of vertical linkages between components. Samples collected from these components were categorized into broiler samples, environment samples, or product samples, which are listed in Table 2. Samples collected from the slaughter facilities were further stratified by sample type and, for broiler product samples, by processing stage, including product samples collected before picking (pre-pick samples), after picking but before evisceration (post-pick samples), after evisceration but before chilling (pre-chill samples), and after chilling (post-chill samples) (Table 2). Subgroups defined by component-sample-type combinations across the included studies were used in further data synthesis and meta-analysis.

**Table 2.** Summary of detailed sample type at each PDN component.

| PDN component | Sample Type (Category) | Description |
|---|---|---|
| Farm | Environmental samples | Feces, soil, boots (or boot covers), litter, nets, bedding, feed, drinking water, worker hand swabs, flypapers, sponge-stick swabs, beetle traps, bedding material of rice husk, drag swabs, surface swabs (floors, and walls) |
|  | Broiler samples | Intestinal contents, cecal samples, fresh fecal swabs, and cloacal swabs for broilers at the end of rearing stage |
| Transportation | Environmental samples | Swabs from animal transport boxes, sponge-stick swabs from transport cages’ floor, drag samples, fecal samples |
| Live bird market | Broiler samples | Cecal contents, cloacal swabs |
|  | Product samples | Chicken carcasses |
|  | Environmental samples | Boot socks or overshoes |
| Slaughter facility | Environmental samples | Feces, boots, drag swabs, personnel hygiene swabs (hands/aprons), process water (stunning, scalding, and chiller water), equipment surface and processing device swabs, food contact surface swabs (working tables, knives, cutting boards, baskets, and buckets), non-food contact surface swabs (walls, grounds, floors) |
|  | Broiler samples | Intestinal contents, cecal samples, fresh fecal swabs, and cloacal swabs collected before, during, or after evisceration. |
|  | Organ samples | Internal organs or abdominal cavity collected during or after evisceration, but before chilling. |
|  | Pre-pick samples | Carcasses, carcass rinses, skin from carcasses collected before post picking, including bleeding and scalding. |
|  | Post-pick samples | Carcasses, carcass rinses, skin from carcasses collected after post picking, but before evisceration. |
|  | Pre-chill samples | Carcasses, carcass rinses, skin from carcasses collected after evisceration but before chilling |
|  | Post-chill samples | Carcasses, carcass rinses, skin from carcasses, or leg or breast fillet of butchered carcass collected after chilling |
|  | Other product sample | Carcasses, chicken meat that didn’t mention detailed sampling stage |
| Retail | Broiler product samples | Carcasses, chicken meat, organs that are packaged and sold to customers |
|  | Environmental samples | Boot socks or overshoes, swabs of chopping boards |

### 2.6. Data synthesis and statistical analysis

For each included publication, data corresponding to each component-sample-type subgroup were treated as a separate dataset, and an individual sample-level prevalence was calculated from each dataset by dividing the number of positive samples by the total number tested. Quantitative synthesis was restricted to subgroups represented by at least two independent studies, while subgroups represented by a single study were summarized descriptively. All prevalence estimates were logit-transformed.

Initial random-effects meta-analysis was first conducted to estimate pooled prevalence and corresponding 95% confidence intervals (CIs) for each PDN component, and further for each component-sample type subgroup, separately for *Campylobacter* spp. and NTS. Between-study heterogeneity was assessed using *τ*^2^ and *I*^2^, with the *I*^2^ values of 25%, 50%, and 75% interpreted as rough guides to low, moderate, and high heterogeneity, respectively (Higgins et al., 2003). To explore potential sources of heterogeneity, univariable meta-regression analyses were conducted separately for each pathogen at the farm, slaughter-facility, and retail levels, using country, broiler PDN type, and study period as study-level moderators. However, this approach did not distinguish between heterogeneity among individual prevalence estimates within publications and heterogeneity between publications.

Given the high heterogeneity and the dependence among multiple prevalence estimates from different sample types and PDN components contributed by the same publication, multilevel meta-regression (three-level random-effects models) were subsequently fitted, accounting for dependence among individual prevalence estimates from different component-sample-type subgroups within the same publication and allowing residual heterogeneity to be estimated separately within and between publications. Subgroup was included in the models as a categorical fixed-effect moderator. The three-level structure comprised the sampling variance of individual prevalence estimates (level 1), residual heterogeneity among subgroup estimates within publications (level 2), and heterogeneity between publications (level 3). Random-effects variance components were estimated using Restricted Maximum Likelihood (REML).

Model-based predicted prevalence estimates, and 95% CIs were derived for each available subgroup. Heterogeneity was quantified using the level-two and three random-effects variances, 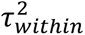 and 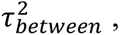, respectively, and their corresponding level-specific *I*^2^ statistics. Differences among PDN component-sample type combinations were assessed through pairwise comparison from these fitted models, using the ‘pwcompare’ command (College Station, TX: Stata Press, 2025). Publication bias assessment via funnel plots and small-study effects assessment using Egger’s regression test, which was developed for comparative data, were not pursued due to the methodological limitations of applying comparative-data tests to proportional meta-analyses (Barker et al., 2021). Leave-one-publication-out sensitivity analyses were conducted by omitting all estimates contributed by each publication in turn and refitting the primary multilevel model to assess the robustness of subgroup-specific pooled prevalence estimates. Analyses included only subgroups with a minimum of three independent publications, ensuring that at least two remained after each omission.

All meta-analyses of the data, forest plots, and leave-one-out sensitivity analysis were conducted using Stata version 19.0 (College Station, TX: Stata Press, 2025). The data, Stata code, and codebook required to reproduce the multilevel meta-regression, pairs comparison, and leave-one-publication-out sensitivity analyses are available on Zenodo (https://doi.org/10.5281/zenodo.23169638).

### 2.7. Quality assessment

Quality assessment of all included studies was conducted using a modified Joanna Briggs Institute (JBI) Critical Appraisal Checklist for Studies Reporting Prevalence Data, where five domains were evaluated, including sample strategy, description of study subjects and setting, coverage of sample type, validity of microbiological identification, and reliability and consistency of measurement (Supplementary Table S5) (Munn et al., n.d.). Two reviewers (CX and SH) independently assessed the included studies for quality assessment. Study quality was assessed by evaluating domains and assigning a score of 1 if the criterion was met and 0 if it was not met or was unclear.

## 3. Results

### 3.1. Study selection

A total of 1,856 records (775 for *Campylobacter* spp. and 1,081 for NTS groups) were initially identified from three databases (Fig. 1). After duplicate removal and title-abstract screening in Covidence, 57 studies (28 for *Campylobacter* spp. and 29 for NTS) were deemed eligible for full-text review. Subsequently, 4 *Campylobacter* studies and 6 NTS studies were excluded for various reasons, the most common of which was a lack of prevalence data, as the studies focused solely on bacterial isolates or conducted targeted sampling of known pathogen-positive broiler flocks.

**Fig 1.**
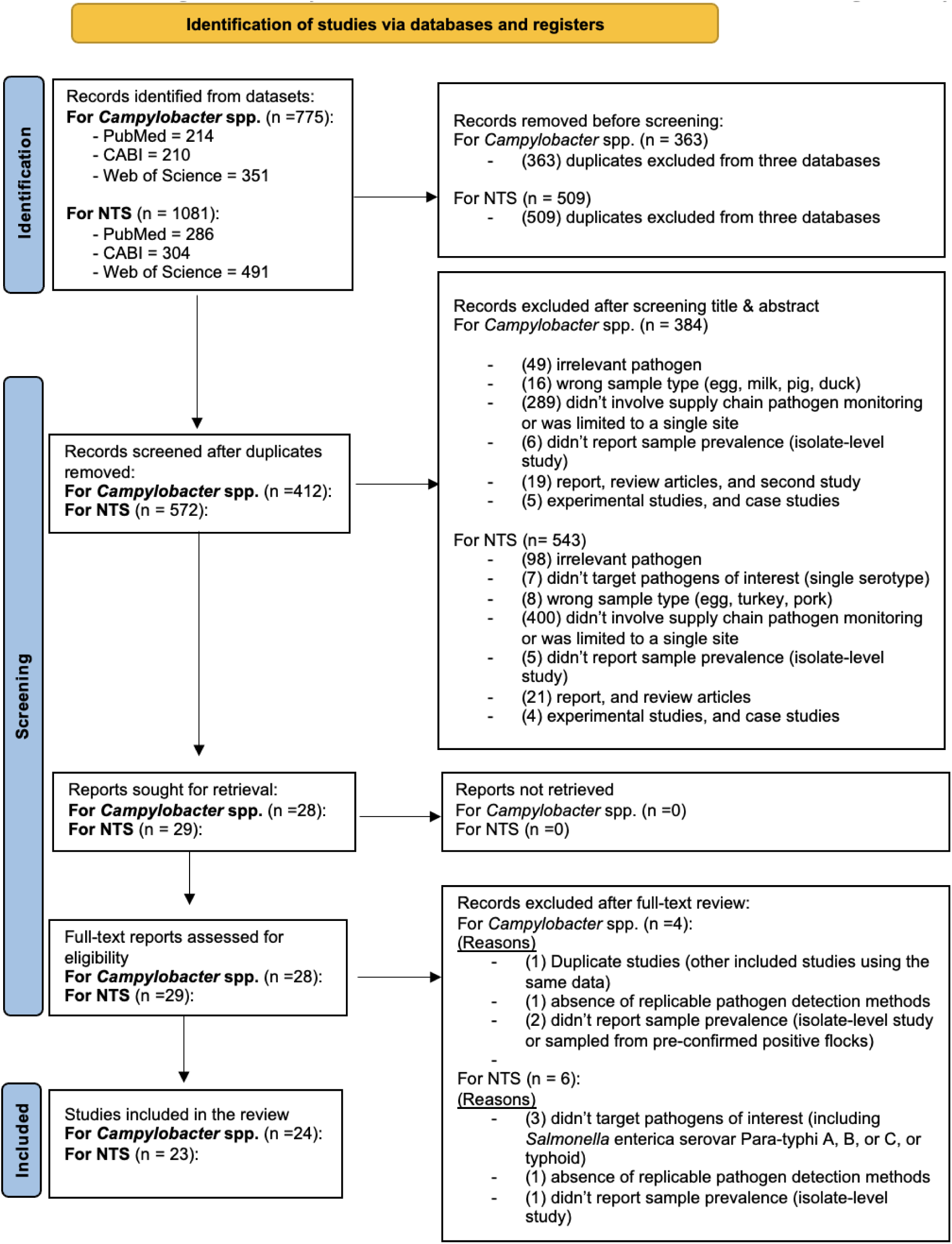
PRISMA flow chart of the systematic review process.

### 3.2. Characteristics of PDN

Broiler PDNs were classified as integrated or non-integrated according to whether sampled production, processing, and distribution components were linked within a vertically integrated system. Integrated PDNs were defined as comprising vertically integrated commercial broiler production and distribution systems, such as integrated broiler companies or complexes, with interconnected sampled components (Fig. 2). In contrast, non-integrated PDNs comprised separately managed systems whose sampled components were not linked within a single vertically integrated network (Fig. 3). Of the studies in the *Campylobacter* spp. group, 12 of 24 (50.0%) were classified as involving integrated PDNs and 12 of 24 (50.0%) as involving non-integrated PDNs. In the NTS group, 11 of 23 studies (47.8%) involved integrated PDNs, whereas 12 of 23 (52.2%) involved non-integrated PDNs.

**Fig. 2.**
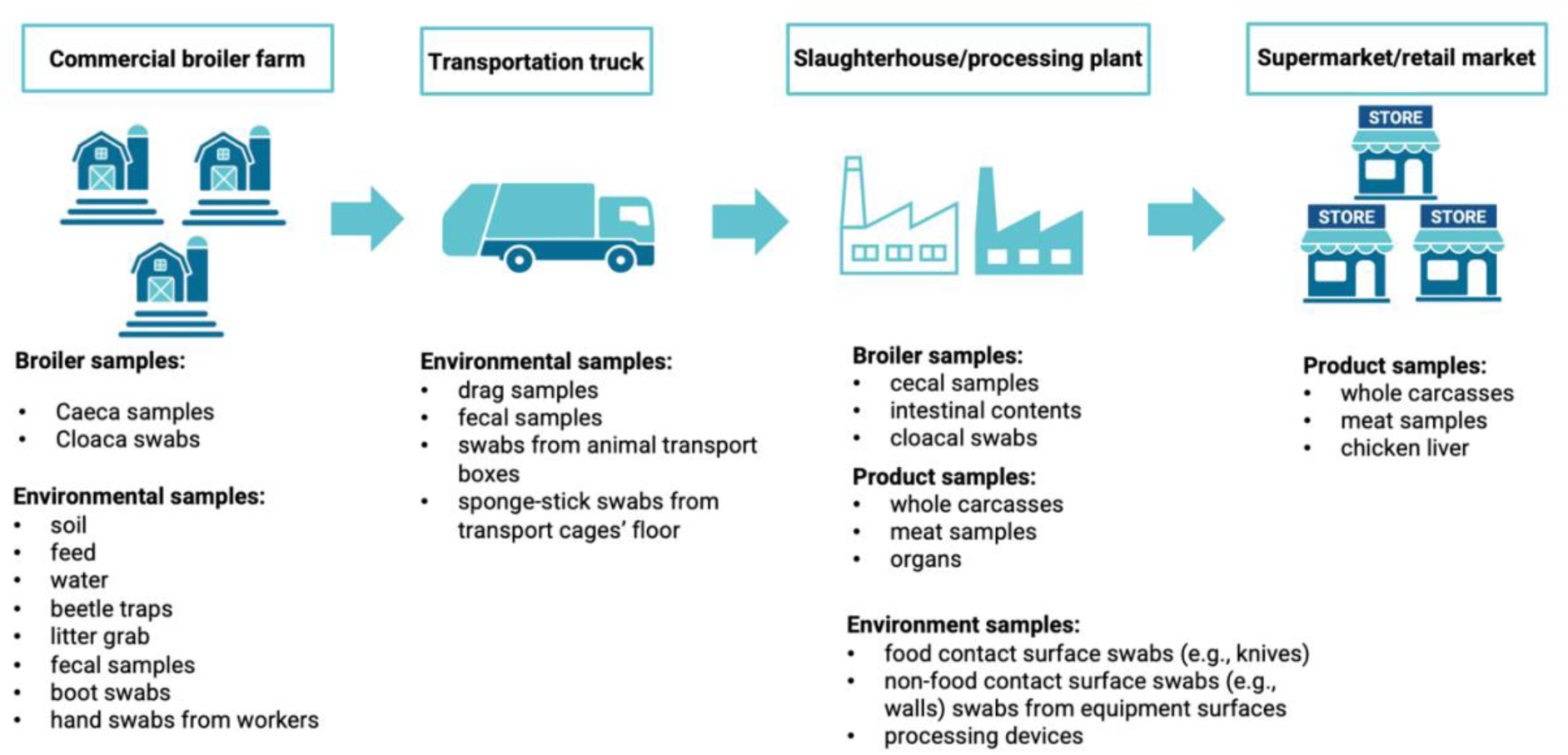
Flow diagram depicting the structure of an integrated PDN and detailing the broiler samples, product samples, and environmental samples collected at different sampling components.

**Fig. 3.**
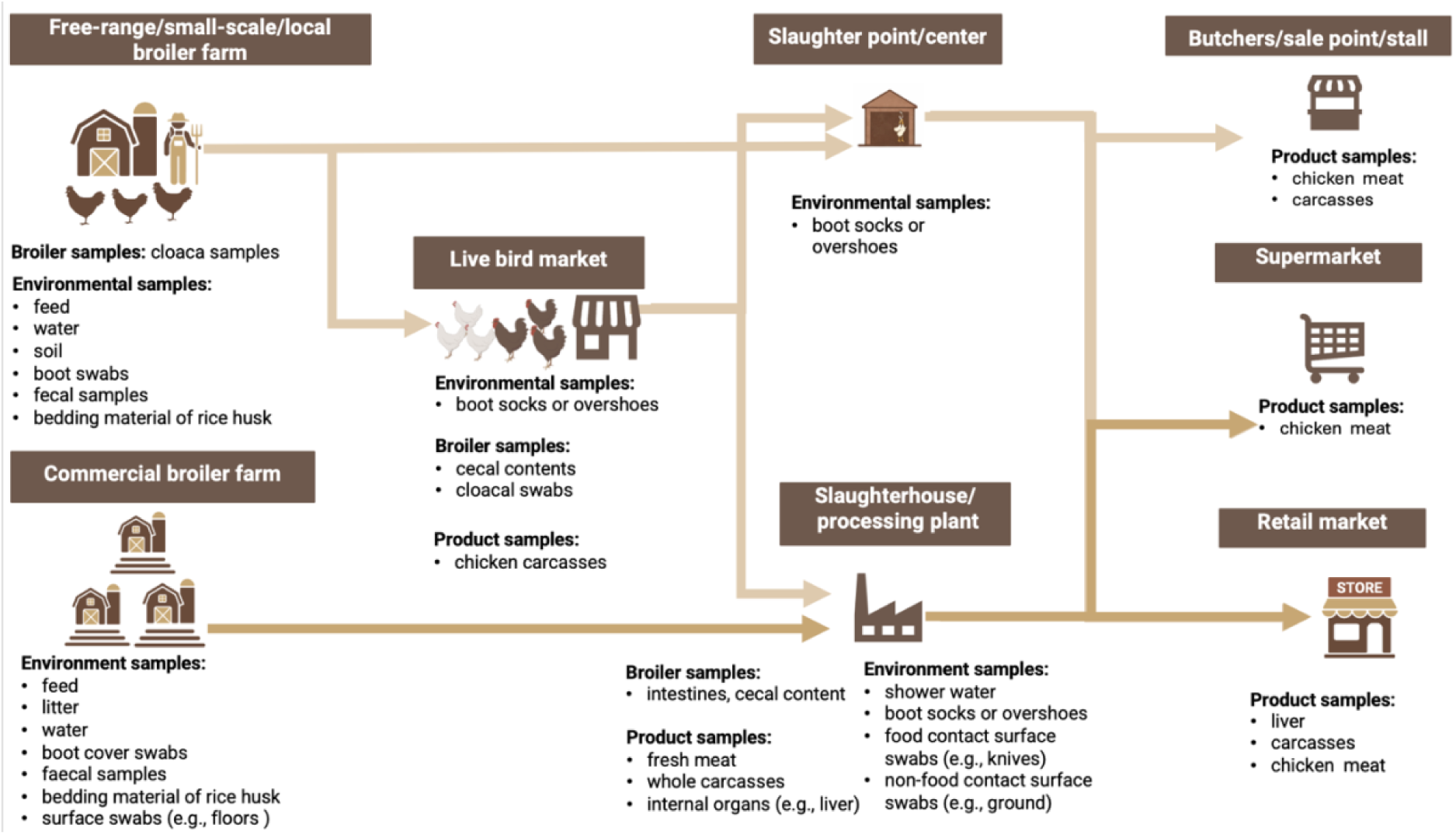
Flow diagram showing the structure of non-integrated PDNs and detailing the broiler samples, product samples and environmental samples collected at different sampling components.

Among the included studies, only those involving integrated PDNs reported follow-up sampling that tracked selected individual birds, flocks or batches from the end of on-farm rearing through transport and slaughter and, in some studies, to retail. Among studies investigating *Campylobacter* spp., (Tang et al., 2020) used plastic identification tags to follow selected broilers from 31 days of age on the farm through slaughter. Similarly, (Iannetti et al., 2020) fitted selected birds in each batch with leg rings before farm departure, and followed up sampling during the slaughtering process. (Adhikari et al., 2024, 2025) conducted follow-up sampling of broiler flocks on transport trucks and subsequently at the processing plant. (M. V. Zbrun et al., 2015) extended batch-level follow-up to retail, sampling broilers from the same flock that were sent together to the same slaughterhouse and sold together at the same retail market. Among studies investigating NTS, only (Adhikari et al., 2024) followed broiler flocks through transport to the processing plant, whereas (Vinueza-Burgos et al., 2019) followed one selected flock per farm to the slaughterhouse. These studies followed pathogen dynamics within the same broiler source along the chain. However, in studies involving non-integrated broiler PDNs, samples from different components were not linked to a common broiler flock, and they originated from different sources. Study settings captured across these components are diverse, encompassing farms of different scales and intensities, live-bird markets, slaughter facilities of different scales (such as slaughterhouses, processing plants, and slaughter points), and retail, such as butchers, sales points, stalls, and supermarkets.

### 3.3. Study-level and subgroup-specific characteristics

For the *Campylobacter* spp. group, a total of 24 included studies were conducted across 15 countries, with Argentina being the most common location (n = 4, 16.7%) (Rossler et al., 2020; M. Zbrun et al., 2013; M. V. Zbrun et al., 2015, 2017). Of these studies, five were conducted during 2000–2010, six during 2011–2015, five during 2016–2020, and three during 2021–2025, while the remaining five did not report their study periods (Supplementary Table S6). For the NTS group, studies (n = 23) were conducted across 12 different countries, most commonly in China (n = 6, 26.1%) (Chen et al., 2019; Cui et al., 2016, 2019; Ma et al., 2017; Ren et al., 2016; Wu et al., 2022). These studies were conducted during 2000–2010 (n = 2), 2011–2015 (n = 7), 2016–2020 (n = 6), and 2021–2025 (n = 5), while the remaining three studies did not report their study periods (Supplementary Table S7). Summaries of the included studies for the *Campylobacter* spp. and NTS groups are provided in Supplementary Tables S6 and S7, respectively.

In the *Campylobacter* spp. group, more than half of studies (62.5%, 15/24) reported pathogen detection in broiler samples from farms or slaughter facilities, including cloacal swabs, cecal samples, intestinal contents, and fresh fecal swabs. Detections in broiler product samples from slaughter facilities or retail, including carcasses, chicken meat, and organs, were reported in 23 studies (95.8%). There were 12 studies (50%) that reported detections in environmental samples collected from farms, transportation trucks, or slaughter facilities, with detailed sample type listed in Table 2. In the NTS group, pathogen detections were reported in broiler samples in 14 of 23 studies (60.9%), product samples in 21 studies (91.3%), and environmental samples in 13 studies (56.5%) across different PDN components.

Subgroup datasets were constructed, stratified by PDN component and sample type. These comprised 90 datasets from 24 articles for the *Campylobacter* spp. group and 82 datasets from 23 articles for the NTS group. For the *Campylobacter* spp. group, approximately half of the datasets (48.9%, 44/90) used selective enrichment followed by molecular identification using Polymerase Chain Reaction (PCR) or Whole Genome Sequencing (WGS). In contrast, only a small number of datasets (18/90, 20.0%) used direct plating followed by DNA-based identification by PCR, primarily from cecal or cloacal samples. For the NTS group, almost one-third of datasets (29.3%, 24/82) used molecular techniques specifically for serotyping, whereas traditional phenotypic serotyping based on the Kauffmann-White scheme remained the majority (56.1%, 46/82).

### 3.3 Persistent heterogeneity across initial meta-analysis, subgroup analyses, and meta-regression

The initial meta-analysis of overall pooled prevalence for each PDN component showed substantial heterogeneity in the farm, slaughter facility, and retail components for both pathogens (all *I*^2^> 96%; Supplementary Figures S1–S11). For *Campylobacter* spp., the value of *I*^2^ exceeded 94% in all component-sample type subgroups within farms and slaughter facilities, except for pre-pick samples at slaughter facilities (*I*^2^= 26.10%; Supplementary Table S8). For NTS, the value of *I*^2^exceeded 92% in all component-sample-type subgroups within farms and slaughter facilities (Supplementary Table S9). The univariable meta-regression analyses further showed residual heterogeneity remained high for *Campylobacter* spp. at the farm (*I*^2^> 99%), slaughter-facility (*I*^2^> 99%) and the retail (*I*^2^> 95%) levels (Supplementary Table S10). For NTS, residual heterogeneity also remained high at the farm (*I*^2^ > 95%), slaughter-facility (*I*^2^> 94%) and retail (*I*^2^> 95%) levels (Supplementary Table S11).

### 3.4. Multilevel meta-regression analysis of *Campylobacter* spp

All selected studies (n = 24), contributing 90 subgroup-specific datasets, were included in the multilevel meta-regression analysis to predict *Campylobacter* spp. prevalence.

On broiler farms, the model-predicted prevalence of *Campylobacter* spp. was significantly lower in environmental samples (16.6%; 95% CI, 8.2–30.9%) than in broiler samples (50.3%; 95% CI, 31.2–69.4%) (OR = 0.197; 95% CI, 0.066–0.592; *p* = 0.004; Fig. 4 and Table 3). At subsequent components along the PDN, the estimated prevalence was 53.9% (95% CI, 14.1–89.3%) in environmental samples during transportation and 22.4% (95% CI, 4.1–65.9%) in broiler samples from live-bird markets. At the slaughter-facility level, no statistically significant differences were detected in pairwise comparisons among sample types (all *p* > 0.05; Table 3). The predicted prevalence in product samples was 49.4% (95% CI, 12.8-86.6%) before picking (pre-pick), 56.8% (95% CI, 28.7-81.1%) after picking but before evisceration (post-pick), 58.9% (95% CI, 29.0-83.4%) after evisceration but before chilling (pre-chill), and 44.8% (95% CI, 25.8-65.6%) after chilling (post-chill). The predicted prevalence in environmental and broiler samples at slaughter facilities was 40.0% (95% CI, 18.5-66.2%) and 50.2% (95% CI, 28.8-71.5%), respectively. At retail, there was a prevalence of 34.9% (95% CI, 20.5-52.7%) among product samples. Residual heterogeneity was greater among subgroups within publications (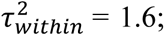 *I*^2^ = 79.7%) than between publications (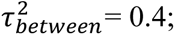 *I*^2^ = 18.9%).

**Fig. 4.**
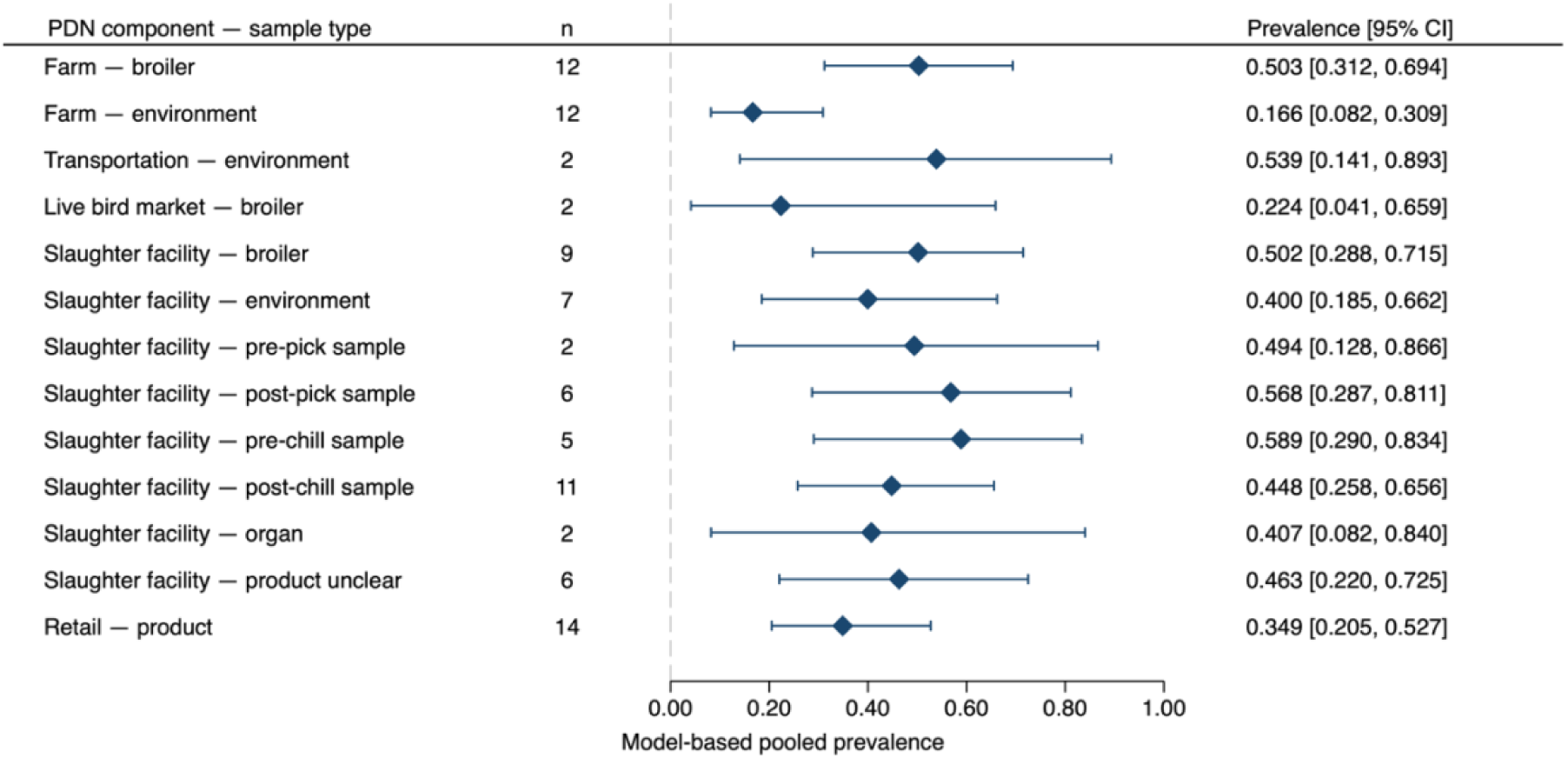
Mean prevalence of *Campylobacter* spp. with 95% CI by PDN component and sample type from multilevel meta-regression. The “n” represents the number of publications contributing to each component-sample-type subgroup prevalence estimate. Diamonds indicate model-based pooled prevalence; whiskers indicate 95% CI.

**Table 3.** Model-based pairwise comparisons of both *Campylobacter* spp. and NTS prevalence among sample types within PDN components.

| **Pathogen** | **Site** | **Comparison** | **OR (95% CI) \*** | ***p* value\*** |
|---|---|---|---|---|
| ***Campylobacter* spp.** | farm | environment vs broiler | 0.197 (0.066 - 0.592) | 0.004 |
|  | slaughter facility | environment vs broiler | 0.661 (0.168 - 2.596) | 0.553 |
|  |  | organ vs broiler | 0.681 (0.768 - 6.051) | 0.731 |
|  |  | post-chill sample vs broiler | 0.806 (0.2441 - 2.666) | 0.725 |
|  |  | post-pick sample vs broiler | 1.305 (0.307 - 5.546) | 0.718 |
|  |  | pre-chill sample vs broiler | 1.421 (0.317 - 6.367) | 0.646 |
|  |  | pre-pick sample vs broiler | 0.969 (0.123 - 7.661) | 0.977 |
|  |  | organ vs environment | 1.030 (0.107 - 9.975) | 0.979 |
|  |  | post-chill sample vs environment | 1.220 (0.331 - 4.502) | 0.765 |
|  |  | post-pick sample vs environment | 1.974 (0.426 - 9.146) | 0.385 |
|  |  | pre-chill sample vs environment | 2.150 (0.435 - 10.620) | 0.347 |
|  |  | pre-pick sample vs environment | 1.466 (0.177 - 12.117) | 0.722 |
|  |  | post-chill sample vs organ | 1.183 (0.136 - 10.279) | 0.879 |
|  |  | post-pick sample vs organ | 1.914 (0.189 - 19.390) | 0.582 |
|  |  | pre-chill sample vs organ | 2.085 (0.199 - 21.816) | 0.539 |
|  |  | pre-pick sample vs organ | 1.422 (0.095 - 21.337) | 0.799 |
|  |  | post-pick sample vs post-chill sample | 1.618 (0.402 - 6.518) | 0.498 |
|  |  | pre-chill sample vs post-chill sample | 1.762 (0.410 - 7.574) | 0.446 |
|  |  | pre-pick sample vs post-chill sample | 1.202 (0.159 - 9.105) | 0.859 |
|  |  | pre-chill sample vs post-chill sample | 1.089 (0.211 - 5.617) | 0.919 |
|  |  | pre-pick sample vs post-chill sample | 0.743 (0.086 - 6.396) | 0.787 |
|  |  | pre-pick sample vs pre-chill sample | 0.682 (0.077 - 6.061) | 0.731 |
| **NTS** | farm | environment vs broiler | 1 (0.498 – 1.991) | 0.991 |
|  | slaughter facility | environment vs broiler | 0.705 (0.296 - 1.676) | 0.429 |
|  |  | organ vs broiler | 0.558 (0.132 - 2.360) | 0.428 |
|  |  | post-chill vs broiler | 0.939 (0.351 - 2.510) | 0.9 |
|  |  | pre-chill vs broiler | 0.702 (0.224 - 2.200) | 0.543 |
|  |  | organ vs environment | 0.792 (0.202 - 3.101) | 0.738 |
|  |  | post-chill vs environment | 1.332 (0.537 - 3.308) | 0.536 |
|  |  | pre-chill vs environment | 0.995 (0.355 - 2.791) | 0.993 |
|  |  | post-chill vs organ | 1.682 (0.378 - 7.479) | 0.495 |
|  |  | pre-chill vs organ | 1.257 (0.287 - 5.495) | 0.762 |
|  |  | pre-chill vs post-chill | 0.747 (0.235 - 2.380) | 0.622 |
|  | retail | product vs environment | 0.832 (0.223 – 3.104) | 0.784 |

\*ORs: The odds ratio and 95% CI were derived from pairwise contrasts in the multilevel random-effects model
\**p* value: Two-sided Wald tests assessed whether the difference in model-estimated logit prevalence between the compared groups was zero (equivalently, OR = 1)

Leave-one-publication-out analyses assessed the sensitivity of predicted prevalence estimates to individual publications across component-sample-type subgroups (Supplementary Figures S11-13). These omissions shifted the point estimates for post-pick and pre-chill samples at slaughter facilities, although their confidence intervals overlapped those of the full models. For post-pick samples, the full-model estimate was 56.8% (95% CI, 28.7-81.1%). This estimate decreased to 44.8% (95% CI, 20.1-72.3%) after omitting (Jozwiak et al., 2006) and increased to 68.5% (95% CI, 36.2-89.3%) after omitting (Tang et al., 2020). For pre-chill samples, the full-model estimate was 58.9% (95% CI, 29.0-83.4%). This estimate decreased to 44.0% (95% CI, 18.5-73.1%) after omitting (Jozwiak et al., 2006) and increased to 70.5% (95% CI, 36.5–90.9%) after omitting (Meeyam et al., 2004). All publications were retained in the primary analyses as they met the eligibility criteria.

### 3.5. Multilevel meta-regression analysis of NTS

The multilevel meta-regression analysis for NTS group included 82 component-sample-type datasets from 23 eligible studies.

On broiler farms, the predicted mean value of NTS prevalence was 17.0% (95% CI, 9.0-29.8%) in environmental samples and 17.1% (95% CI, 8.8-30.5%) in broiler samples, with no significant difference between them (*p* > 0.05, Fig. 5, Table 4). At downstream components of broiler PDNs, mean prevalence was 49.6% during transportation (95% CI, 14.3-85.3%). At live-bird markets, only one contributing dataset was available for each sample type, yielding prevalence estimates of 54.5% for environmental samples and 77.8% for product samples. At the slaughter-facility level, the broiler samples (34.4%; 95% CI, 17.6-56.2%) showed significantly higher prevalence compared to broiler samples collected in farms (OR = 2.54; 95% CI, 1.058-6.109; *p* = 0.037, Table 4). However, no significant differences appeared among the subgroups in the pairwise comparisons (*p* > 0.05) (Table 3). The predicted NTS prevalence was 27% (95% CI, 14.8-44.1%) in environmental samples. Product samples collected before chilling (pre-chilling) and after chilling (post-chill) had the predicted mean prevalence of 26.9% (95% CI, 11.4-51.3%) and 33% (95% CI, 16.2-55.6%), respectively. Product samples collected at the retail stage had a predicted prevalence of 27.1% (95% CI, 15.9-42.2%) and 30.9% (95% CI, 10.2-63.8%) for environmental samples. Residual heterogeneity was greater between publications (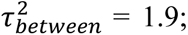 *I*^2^ = 80.3%) than between component-sample-type subgroups within publications (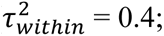 *I*^2^ = 17.4%).

**Fig. 5.**
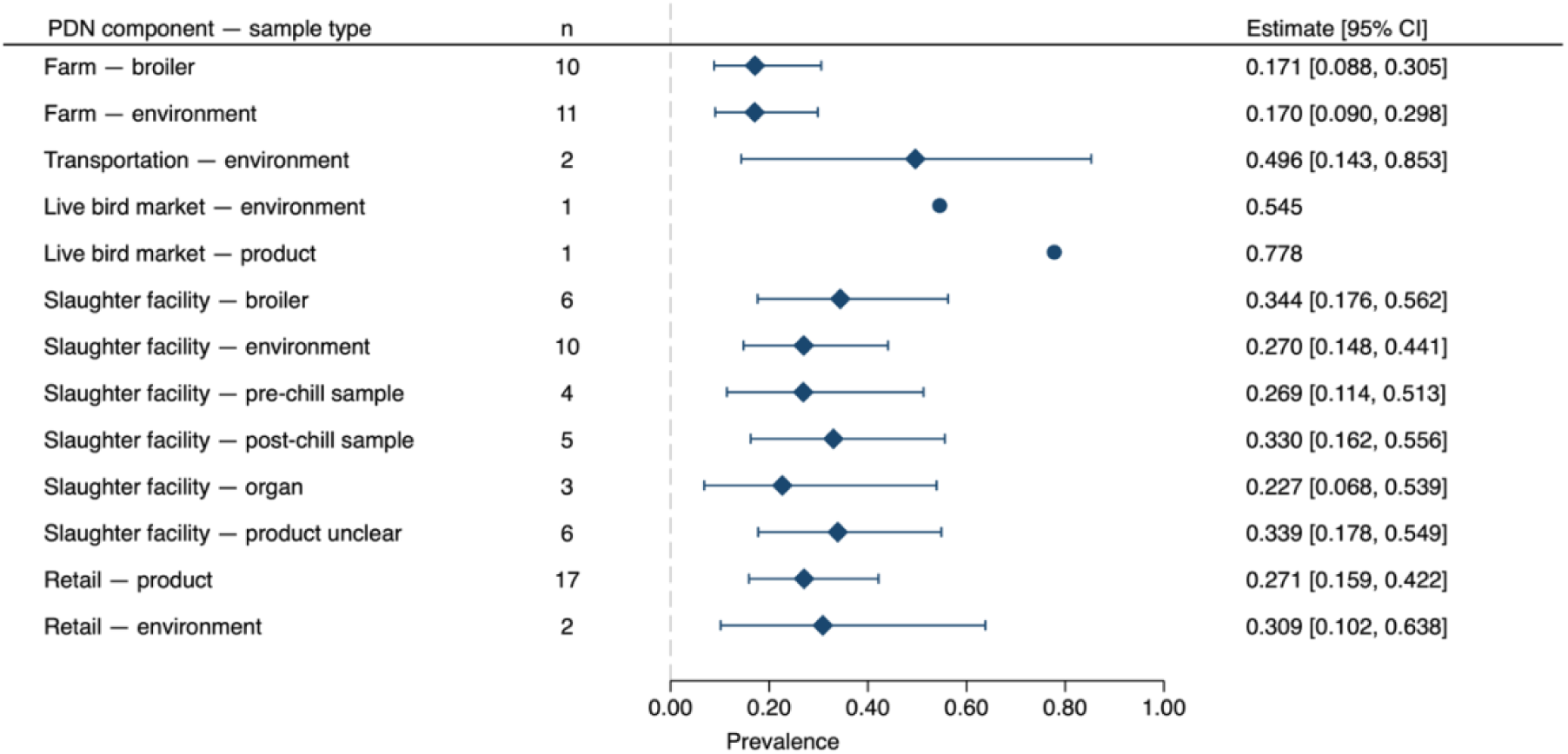
Mean prevalence of NTS with 95% CI by PDN component and sample type from multilevel random-effects models. The “n” represents the number of publications contributing to each component-sample-type subgroup prevalence estimate. Diamonds indicate model-based pooled prevalence; whiskers indicate 95% CI. Circles are raw observed prevalences for the transportation singleton group, with no CI shown.

**Table 4.** Model-based pairwise comparisons of both *Campylobacter* spp. and NTS prevalence among PDN components for the same sample type.

| **Pathogen** | **Sample type** | **PDN-component comparison** | **OR (95% CI) \*** | ***p* value\*** |
|---|---|---|---|---|
| *Campylobacter* spp. | broiler | live bird market vs farm | 0.283 (0.038-2.120) | 0.220 |
|  |  | slaughter facility vs farm | 0.994 (0.312 - 3.157) | 0.992 |
|  |  | slaughter facility vs live bird market | 3.496 (0.441 - 27.718) | 0.441 |
|  | environment | transportation vs farm | 5.851 (0.732 - 46.776) | 0.096 |
|  |  | slaughter facility vs farm | 3.334 (0.937 - 11.866) | 0.063 |
|  |  | transportation vs slaughter facility | 1.755 (0.198 - 15.539) | 0.613 |
| NTS | broiler | slaughter facility vs farm | 2.54 (1.058 - 6.109) | 0.037 |
|  | environment | retail vs farm | 2.172 (0.593- 7.959) | 0.242 |
|  |  | slaughter facility vs farm | 1.799 (0.957- 3.381) | 0.068 |
|  |  | transportation vs farm | 4.793 (0.868 - 26.459) | 0.072 |
|  |  | slaughter facility vs retail | 0.829 (0.227- 3.028) | 0.776 |
|  |  | transportation vs retail | 2.207 (0.272 - 17.934) | 0.459 |
|  |  | transportation vs slaughter facility | 2.664 (0.474 -14.972) | 0.266 |

\*ORs: The odds ratio and 95% CI were derived from pairwise contrasts in the multilevel random-effects model
\**p* value: Two-sided Wald tests assessed whether the difference in model-estimated logit prevalence between the compared groups was zero (equivalently, OR = 1)

In leave-one-publication-out analyses, omitting (Bellil et al., 2025) shifted the predicted prevalence estimate for organ samples at slaughter facilities, where the estimate increased from 22.7% (95% CI, 6.8-53.9%) in the full model to 34.6% (95% CI, 9.9-71.7%) after omission (Supplementary Figures S15-17). All studies were retained in the primary analyses as they met the eligibility criteria.

### 3.6. Diversity of NTS serotypes

In total, 66 NTS serovars were identified among 1,593 isolates from samples collected at farms, slaughter facilities, and retail, as reported in 17 publications from 11 countries (Fig. 6A and 6B). The 10 most frequently reported broiler chicken-associated serotypes were Infantis, Enteritidis, Kentucky, Hadar, Typhimurium, Derby, Weltevreden, Indiana, Meleagridis, and Agona. The frequency and diversity of NTS serotypes varied across components. Infantis was the most frequently reported serotype in retail (31.0%), whereas Enteritidis was the most frequently reported in farms (20.2%) and slaughter facilities (19.3%) (Fig. 6A, Supplementary Table S 12). Retail and slaughter facilities reported the greatest diversity of serotypes (44 different serotypes), followed by farms (33 different serotypes). Serotype frequency and diversity also differed among sample types. Enteritidis predominated in broiler samples from farms (30.4%, 41/135), whereas Hadar was the most common in environmental samples from farms (22.0%, 37/168) and slaughter facilities (38.9%, 37/95) (Fig. 6C-D), and Infantis was the most frequently reported serotype in retail product samples (31.6%, 213/674).

**Fig. 6.**
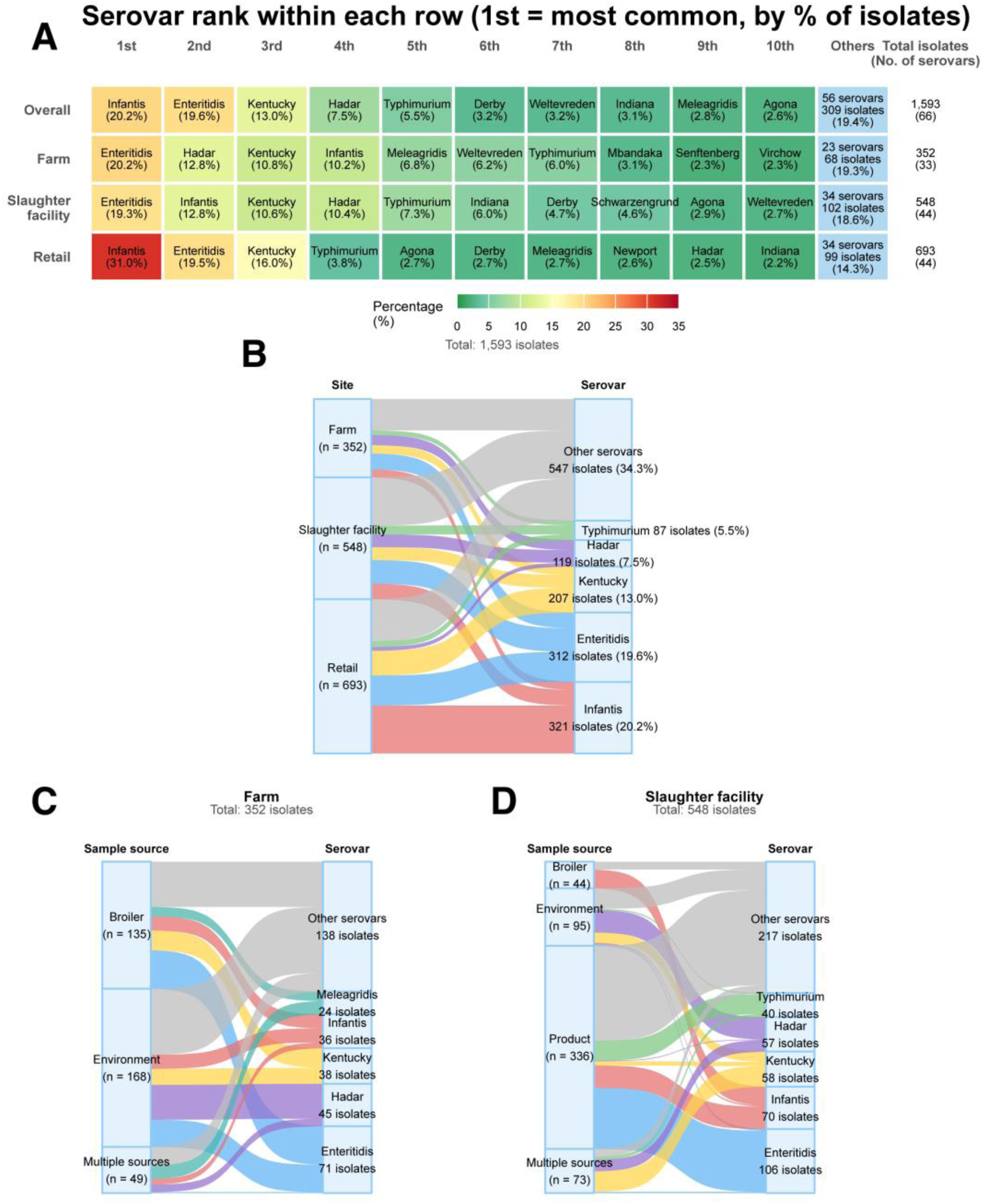
Distribution of NTS serovars across PDN components and sample sources. (A) Heatmap showing the 10 most frequently reported serovars overall and within farms, slaughter facilities, and retail. (B–D) Sankey diagrams showing serovar distributions across PDN components (B) and by sample source within farms (C) and slaughter facilities (D). Ribbon widths are proportional to isolate counts, with the total flow in each panel normalized to 100%. “Other serovars” groups serovars not displayed individually in the Sankey diagrams. “Multiple sources” denotes isolates reported collectively across two or more sample-source categories for which source-specific counts were unavailable.

### 3.7. Quality assessment

Supplementary Tables S 13 and 14 summarize the quality assessment using the modified JBI checklist. Among the 24 *Campylobacter* spp. studies, 15 (62.5%) met the criteria for sampling strategies at both site and sample levels. Study subjects, settings, and sampling procedures were described in detail in 19 studies (79.2%). All 24 studies (100.0%) achieved coverage across sample types and components, with analyzed sample numbers closely matching those planned or collected and no substantial unexplained losses or exclusions. Valid microbiological identification was reported in 20 studies (83.3%), using standard isolation and identification methods, confirmatory testing, and species-level identification. Reliability and consistency of measurement were reported in 18 studies (75.0%), with standardized sample handling and laboratory procedures.

Among the 23 NTS studies, six (26.1%) met the sampling strategies criteria, and 17 (73.9%) provided detailed descriptions of study subjects, settings, and sampling procedures. Adequate coverage across sample types and sites was achieved by 22 studies (95.7%). Valid microbiological identification, including serovar determination by standardized serotyping or validated molecular methods, was reported in 18 studies (78.3%). Reliable and consistent measurement was reported in 22 studies (95.7%).

## 4. Discussion

This systematic review provides, to our knowledge, the first integrated meta-analysis of *Campylobacter* spp. and NTS prevalence across the broiler PDN from farm to retail. By pooling data from 24 *Campylobacter* studies and 23 NTS studies across 15 and 12 countries, respectively, we find that slaughter facilities are critical contamination hotspots for both pathogens, with the predicted prevalence of *Campylobacter* spp. reaching 58.9% in samples collected from products, especially after evisceration but before chilling (pre-chill), and NTS of 34.4% in broiler samples collected before, during, or after evisceration. The reported detection of pathogens at other post-farm stages, including transportation, live bird market, and retail, also highlights the need for stage-specific risk assessment and targeted interventions.

Our review first maps the structure of broiler PDNs and identifies where pathogens have been detected, considering the movement of broiler chickens and their products through the supply chain in both integrated and non-integrated systems. Integrated PDNs generally included commercial farms, transportation trucks, slaughterhouses, processing plants, and retail. Non-integrated PDNs varied across countries and studies and included main PDN components, such as farms, slaughter, retail, all operating at different scales, as well as intermediate nodes such as live-bird markets. Broiler movements through these networks create multiple opportunities for pathogen contamination of broiler chickens, their products, and the surrounding environment. At the farm level, broiler chickens serve as reservoirs for *Campylobacter* and *Salmonella*, which colonize the gastrointestinal tract (Rasschaert, Houf, Van Hende, et al., 2007). Once broilers become colonized, they may shed the pathogens in their feces, contaminating litter and the surrounding farm environment and potentially spreading contamination to soil, water, and feed (Heyndrickx et al., 2002; Newell & Fearnley, 2003). This fecal contamination, which contains these pathogens, can also facilitate rapid fecal-oral transmission within the flock (Neelawala et al., 2024; Zainol et al., 2024). Further cross-contamination to non-infected broiler chickens may occur during transport to slaughter through contact with fecal material and contaminated transport crates, which may contribute to downstream carcass and product contamination. During slaughter and subsequent processing operations, including defeathering, evisceration, chilling, and carcass handling, broiler carcasses and products may become contaminated with these pathogens through contact with fecal material or intestinal contents originating from the same flock or from previously slaughtered flocks (Zeng et al., 2021). Retail is another potential component of consumer exposure, as contaminated chicken products and their packaging may introduce pathogens into the retail environment and facilitate cross-contamination (Harrison et al., 2001).

*Campylobacter* spp. contamination was common in product samples at several stages of slaughter processing, with a predicted overall prevalence of 44.8% (95% CI, 25.8–65.6%) even after chilling. Previous studies also provided evidence of *Campylobacter* contamination at different stages during processing. A systematic review examined changes in *Campylobacter* prevalence on chicken carcasses during processing and found that defeathering and evisceration can increase *Campylobacter* contamination, whereas chilling often reduces contamination levels (Guerin et al., 2010). Another study analyzed *Campylobacter* prevalence at different processing steps in poultry slaughterhouses and found that sampling points with the highest contamination rates were after evisceration and decreased significantly after chilling (Lee et al., 2017). The higher predicted prevalence of *Campylobacter* spp. after evisceration may partly reflect contamination from intestinal contents released during viscera removal (Boysen & Rosenquist, 2009). Contaminated processing equipment may provide an additional route for bacterial transfer between carcasses, contributing to cross-contamination during processing (Posch et al., 2006). Evidence from chilling and post-chilling interventions indicates that *Campylobacter* contamination can be reduced, although effectiveness varies with the treatment and outcome measured (Leone et al., 2024). A meta-analysis covering 21 European countries also reported lower *Campylobacter* prevalence in frozen than chilled poultry meat (Gonçalves-Tenório et al., 2018). Therefore, our prevalence estimates and these process-stage findings support monitoring these pathogens at multiple slaughter stages and implementing measures to prevent intestinal leakage, limit equipment-mediated cross-contamination, and ensure effective chilling.

Similar variation in the predicted NTS prevalence was also found across PDN components and sample types. In broiler samples, predicted prevalence at slaughter facilities (34.4%; 95% CI, 17.6–56.2%) was approximately twice that on farms at the end of rearing (17.1%; 95% CI, 8.8–30.5%). This difference may partly reflect exposure to contaminated environments during transport, consistent with the high predicted NTS prevalence in transport-associated environmental samples (49.6%; 95% CI, 14.3–85.3%). Higher *Salmonella* detection in fecal samples after transport to the slaughterhouse has also been reported (Marin & Lainez, 2009). During live broiler transport, transport stress may increase fecal shedding by colonized birds, enabling these pathogens to contaminate transport crates and broiler surfaces, facilitate cross-contamination among birds, and be carried into slaughter facilities, where they may contribute to downstream carcass and product contamination (Mulder, 1995; Slader et al., 2002). Meanwhile, transport crates can remain contaminated after washing and disinfection, creating a potential exposure source for broilers (Corry et al., 2002). Therefore, transport hygiene warrants greater attention in NTS control across the PDN. Given the limited evidence base, longitudinal studies tracking the same flocks and their transport environments from farms to slaughter are needed to help quantify the contribution of transport-associated exposure to downstream NTS contamination.

Our review identified Infantis, Enteritidis, Kentucky, Hadar, and Typhimurium as the top five serovars among all included studies. The predominant serovar varied by PDN component and sample type, with Enteritidis most common in broiler samples from farms, Hadar in environmental samples from farms and slaughter facilities, and Infantis in retail product samples. These findings suggest that sampling on different types, such as broiler, environmental, and product, could provide complementary information for NTS surveillance across broiler PDNs, to better identify the predominant serovars in different sample types and guide the selection of PDN components and sample types for subsequent surveillance and help prioritize serovars for further investigation. Integrating these sampling approaches with serotyping could therefore help characterize serovar distributions across production, processing, and retail.

Enteritidis is a major cause of human salmonellosis worldwide. International surveillance during 2001–2007 identified Enteritidis as the most frequently reported serotype among human *Salmonella* isolates in Asia, Latin America, Africa, and Europe (Hendriksen et al., 2011). A United States genomic source-attribution analysis identified chicken as the most common predicted source of human Enteritidis infections (Rose et al., 2025). European Union surveillance data for 2017 showed that broiler flocks and chicken meat accounted for 57.2% of the Enteritidis isolates reported from the food and animal sources (European Food Safety Authority and European Centre for Disease Prevention and Control (EFSA and ECDC), 2018). These findings support targeted monitoring and control of Enteritidis during broiler production.

Hadar was among the top three serovars in Europe associated with human *Salmonella* infections during 2001 to 2007 (Hendriksen et al., 2011). A recent analysis of the United States pangenome indicated that a Hadar lineage, which became dominant in 2020, originated from commercial poultry populations and later spread to backyard poultry environments, emphasizing the circulation of contamination across poultry systems (Tagg et al., 2026). Together with our environmental findings, this supports closer investigation of Hadar transmission within poultry production and processing environments which can serve as critical reservoirs for this pathogen.

A high burden of illness caused by Infantis in chicken-associated outbreak illnesses during 2017–2021 was reported in an analysis of CDC’s Foodborne Disease Outbreak Surveillance System (FDOSS) data (Marshall et al., 2024). In addition, the detection of genomically related Infantis isolates in retail poultry products over multiple months from a study conducted in Netherlands suggests that related strains may persist in the poultry production chain, potentially with a contribution from the pESI megaplasmid (van der Voort et al., 2026). The prominence of Infantis in retail samples, together with its documented contribution to chicken-associated outbreak illnesses in humans and its potential persistence in the production chain, supports targeted surveillance and control of Infantis in chicken products.

Several limitations of this systematic review and meta-analysis should be acknowledged. First, although funnel plots and statistical tests, such as Egger’s and Begg’s tests, are commonly used to assess publication bias (Begg & Mazumdar, 1994; Egger et al., 1997), these methods were originally developed for comparative studies and rely on the assumption that studies with positive results are published more often than those with negative results (Barker et al., 2021). Since our meta-analysis synthesized single-arm proportions without comparative groups, the concept of a positive result does not apply. Therefore, these statistical tests were omitted to avoid misleading findings. To assess the methodological quality of the primary studies, a qualitative appraisal was conducted for all included studies using a modified JBI Critical Appraisal Checklist for Studies Reporting Prevalence Data (Munn et al., n.d.), which covers five domains. Given that individual domains carry different methodological weights, calculating a total score for each study across five domains was not chosen; instead, a domain-specific qualitative summary was conducted to provide a more comprehensive evaluation across all included studies. Third, studies sampling multiple PDN components were included, and predicted prevalence estimates across component-sample-type subgroups were compared. However, the same source of broilers was followed across stages in only four studies of *Campylobacter* spp. and two studies of NTS. Differences in predicted prevalence should therefore not be interpreted as within-same-broiler-source changes. Finally, the initial conventional meta-analysis showed substantial heterogeneity at the PDN component (*I*^2^ > 96%) and component-sample type subgroup (*I*^2^ > 92%) levels for both *Campylobacter* spp. and NTS. In the following meta-regression, substantial residual heterogeneity (*I*^2^> 94%) still persisted after adjustment for study-level moderators, including country, production system (integrated versus non-integrated), and study period. However, this approach did not distinguish between heterogeneity among individual prevalence estimates within publications and heterogeneity between publications. We therefore fitted a multilevel meta-regression analysis, a three-level random-effects model, which allowed residual heterogeneity to be quantified separately within and between publications while accounting for the dependence among estimates.

In conclusion, slaughter facilities showed high prevalence of *Campylobacter* spp. and NTS, supporting their prioritization for targeted monitoring and implementation of contamination-control measures. Infantis, Enteritidis, Kentucky, Hadar, and Typhimurium were the five most common NTS serovars across the included studies. Given the observed variation in NTS serovar distribution, surveillance should include multiple sample types across network components. Our model-based estimates of pathogen prevalence provide baseline inputs for future risk assessments of foodborne pathogen transmission along the broiler production and distribution network.

## CRediT authorship contribution statement

Chen Xin: Writing - review & editing, Writing - original draft, Conceptualization, Data curation, Investigation, Methodology, Software, Formal analysis, Validation, Visualization. Shanshan He: Writing - review & editing, Investigation, and Methodology. Nicola Di Girolamo: Writing - review & editing, Conceptualization, Methodology. Casey L. Cazer: Writing - review & editing, Conceptualization, Methodology, Formal analysis, Supervision, Resources. Priscilla F. Gerber: Writing - review & editing, Conceptualization, Methodology, Formal analysis, Supervision, Resources.

## Acknowledgments

The authors acknowledge the contributions of Christian J. Miller, the librarian at Cornell University, for supporting protocol development and reviewing the electronic search strategy.

## Data availability Statement

The data, Stata code, and documentation details required to reproduce the multilevel meta-regression, pairwise comparisons, and leave-one-publication-out sensitivity analyses are publicly available on Zenodo at: https://doi.org/10.5281/zenodo.23169638. The Cornell Center for Social Sciences verified that the data and replication code replicate the numerical results reported in this article.

