## Supplementary Material for "A systematic review and meta-analysis of the prevalence of *Campylobacter* and non-typhoidal *Salmonella* along broiler chicken production and distribution networks"

### Supplementary Information Tables of Contents

**Supplementary Table S 1:** Keywords and search strings in PubMed

| Keywords | Keyword searched |
|---|---|
| Population | |
| Broiler | broiler* OR chicken* OR “chicken meat”<br><br>broiler*[Title/Abstract] OR chicken*[Title/Abstract] OR "chicken meat"[Title/Abstract] |
| Pathogens | |
| *Campylobacter* | campylobacter* OR campylobacter jejuni OR campylobacter coli OR “C. coli” OR “C. jejuni”<br><br>campylobacter*[Title/Abstract] OR campylobacter jejuni[Title/Abstract] OR campylobacter coli[Title/Abstract] OR "C. coli"[Title/Abstract] OR "C. jejuni"[Title/Abstract] |
| non-typhoidal *Salmonella* | Salmone* OR “nontyphoidal Salmonella” OR “non-typhoidal Salmonella” OR “NTS”<br><br>("salmone*"[Title/Abstract] OR "nontyphoidal Salmonella"[Title/Abstract] OR "non-typhoidal Salmonella"[Title/Abstract] OR "NTS"[Title/Abstract]) |
| Poultry production and distribution network | |
| Chain | “production and distribution network” OR “food chain” OR “supply chain” OR “distribution chain” OR “production chain” OR “value chain” OR “poultry chain” OR (farm* AND market* AND slaughter*)<br><br>"production and distribution network"[Title/Abstract] OR "food chain"[Title/Abstract] OR "supply chain"[Title/Abstract] OR "distribution chain"[Title/Abstract] OR "production chain"[Title/Abstract] OR "value chain"[Title/Abstract] OR "poultry chain"[Title/Abstract] OR (farm*[Title/Abstract] AND market*[Title/Abstract] AND slaughter*[Title/Abstract]) |

- #Broiler AND *Campylobacter* AND Chain
- #Broiler AND non-typhoidal *Salmonella* AND Chain

Filters applied: database inception - October 1$^{st}$, 2025

**Supplementary Table S 2:** Keywords and search strings in CABI

| Keywords | Keyword searched |
|---|---|
| Population | |
| Broiler | broiler* OR chicken* OR “chicken meat”<br><br>ab:(broiler* OR chicken* OR "chicken meat") |
| Pathogens | |
| *Campylobacter* | “Campylobacter” OR "Campylobacter coli" OR "Campylobacter jejuni"<br><br>indexingterm:(Campylobacter OR "Campylobacter coli" OR "Campylobacter jejuni") |
| non-typhoidal *Salmonella* | Salmone* OR “nontyphoidal Salmonella” OR “non-typhoidal Salmonella” OR “NTS”<br><br>ab:(Salmone* OR "nontyphoidal Salmonella" OR "non-typhoidal Salmonella" OR "NTS") |
| Poultry production and distribution network | |
| Chain | “production and distribution network” OR “food chain” OR “supply chain” OR “distribution chain” OR “production chain” OR “value chain” OR “poultry chain”<br><br>ab:("production and distribution network" OR "food chain" OR "supply chain" OR "distribution chain" OR "production chain" OR "value chain" OR "poultry chain") |

- #Broiler AND *Campylobacter* AND Chain
- #Broiler AND non-typhoidal *Salmonella* AND Chain

Filters applied: database inception - October 1st, 2025

**Supplementary Table S 3:** Keywords and search strings in Web of Science

| Keywords | Keyword searched |
|---|---|
| Population | |
| Broiler | TS = (broiler* OR chicken* OR “chicken meat”) |
| Pathogens | |
| *Campylobacter* | TS = (campylobacter* OR “campylobacter jejuni” OR “campylobacter coli” OR “C. coli” OR “C. jejuni”) |
| non-typhoidal *Salmonella* | TS = (Salmone* OR “nontyphoidal Salmonella” OR “non-typhoidal Salmonella” OR “NTS”) |
| Poultry production and distribution network | |
| Chain | TS = (“production and distribution network” OR “food chain” OR “supply chain” OR “distribution chain” OR “production chain” OR “value chain” OR “poultry chain” OR (farm* AND market* AND slaughter*)) |

- #Broiler AND *Campylobacter* AND Chain
- #Broiler AND non-typhoidal *Salmonella* AND Chain

Filters applied: database inception - October 1$^{st}$, 2025

**Supplementary Table S 4:** Data extraction form

| Topic | Description |
|---|---|
| Paper information | • First author's name<br>• Title of the article<br>• Journal<br>• Year of publication |
| Study information | • Design: cross-sectional and case-control<br>• Country of study conducted<br>• Year (Duration) of study conducted (sampling period): (YYYY. MM) - (YYYY. MM) |
| Sampling information | • Target foodborne pathogens: *Campylobacter* spp., NTS<br>• PDN type: integrated PDN/ non-integrated PDN<br>• Sampling strategy:<br>1) non-integrated PDNs: probability /non-probability sampling<br>2) integrated PDN: follow the same batch or not<br>• Sampling components (source): farm, market, slaughter facility<br>• Operational scale of the sampled component [small-scale or large-scale] |
| Sampling and pathogen detection | *(For each sampling component, per sample type, consider extracting the following information)*<br><br>• Sample type:<br>1) broilers: cecal and cloacal samples<br>2) broiler products: meat, carcass, skin, feet, feathers, or other product samples<br>3) environmental samples: feces, litter, water, soil, etc.<br><br>• Sample size per sample type:<br>1) Total number of samples collected or tested (n)<br>2) Positive number (based on descriptive results, raw data, without any simulation from the model): The number of samples that tested positive (n)<br><br>• Sample levels: individual/pool<br><br>• Detection method—enrichment or not (if with enrichment, extract enrichment medium)<br><br>• Detection method-- diagnostic technique<br>1) bacteriological culture<br>2) biochemical test<br>3) molecular (including, WGS, PCR, real-time PCR, multiplex PCR)<br>4) immunological assay (MALDI-TOF)<br>5) for NTS serotype identification: standardized serotyping scheme (e.g., the Kauffmann-White-Le Minor scheme) |

| | |
|---|---|
| Evaluation | *(Evaluation was based on per sample type within each PDN component)*<br><br>• No of positive sample among the total amount per sample type<br>• For NTS serotype: the number of isolates of each serotype (n) |

**Supplementary Table S 5:** Description of methodological quality assessment for included studies

| Domain | Question and Criterion |
|---|---|
| Sample strategy | *Was the sample frame appropriate to address the target population?*<br><br>(1): For integrated PDN:<br>- site level: studies that systematically tracked the same batch/flock across multiple sites.<br>- sample level: samples of each type were randomly selected per site.<br>(1): For non-integrated PDNs:<br>- site level: sites were selected using random, systematic, or population-based approaches.<br>- sample level: samples of each type were randomly selected per site.<br><br>(0): Studies relied on convenience sampling, or sampled isolated sites without a clear, systematic approach. |
| Description of study subjects and setting | *Were the study subjects and the setting described in detail?*<br><br>(1): Clear and detailed description of study subjects and the setting, including:<br>- the geographic location of study conducted<br>- key characteristics of sampling sites (e.g., big or small-scale slaughter facility, commercial or free-range farm)<br>- key characteristics of samples and sampling process<br> - type of sample<br> - detailed sampling period<br> - sample handling, storage, and transportation<br><br>(0): Insufficient description. |
| Coverage of sample type | *Was the data analysis conducted with sufficient coverage of the identified sample?*<br><br>(1) For each sample type at each site, the number of samples included in the analysis is close to the number originally planned or collected, with no substantial unexplained exclusions or missing samples.<br><br>(0) One or more planned sample subgroups (i.e., a specific sample type at a particular site) were substantially lost or excluded from the analysis without appropriate statistical adjustment. Alternatively, the original sampling plan or the extent of sample loss/dropout was not reported. |

| | |
|---|---|
| Validity of microbiological identification (*Campylobacter* spp.) | *Were valid methods used for the identification of the condition?*<br><br>(1) For every sample type at every study site, it should follow the following requirements:<br><br>- Pathogens were isolated and identified using recognized standard methods (e.g., standard culture methods)<br>- Pathogen identification was confirmed using an appropriate method, such as molecular testing (e.g., PCR, real-time PCR, multiplex PCR, or whole-genome sequencing) or MALDI-TOF mass spectrometry.<br>- *Campylobacter* spp. isolates were identified to the species level.<br><br>(0): The study relied solely on presumptive identification (e.g., visual inspection on selective agar) without confirmatory tests.<br>(0): Isolates were reported only at genus level without species-level identification. |
| Validity of microbiological identification (NTS) | *Were valid methods used for the identification of the condition?*<br><br>(1) For every sample type at every study site, it should follow the following requirements:<br><br>- Pathogens were isolated and identified using recognized standard methods (e.g., standard culture methods)<br>- Serovar determination using a standardized serotyping scheme (e.g., the Kauffmann–White–Le Minor scheme) or another clearly defined and validated molecular typing method.<br><br>(0): The study relied solely on presumptive identification (e.g., visual inspection on selective agar).<br>(0) Without serovar determination. |
| Reliability and consistency of measurement | *Was the condition measured in a standard, reliable way for all participants?*<br><br>(1): The study clearly describes a standardized process applied to all samples, including consistent transport/storage conditions and the exact same laboratory methods (or recognized international standard protocols) used across different sites.<br>- If different laboratory methods are required for animal versus environmental samples, a detailed description is provided. |

| | |
|---|---|
| | (0): The study failed to report critical sample handling details, or used inconsistent methodologies for different subsets of samples without clear justification. |

- Note: #1 = criterion met; 0 = criterion not met or uncle

**Supplementary Table S 6:** Summary of study characteristics of all included studies in the *Campylobacter* spp. group (dataset: component-sample type)

| Study | Study year | Study year category | Country | PDN type | Sampling site | PDN component category | Sample source | Sample type category | Detection method | No of positive samples | Sample size |
|---|---|---|---|---|---|---|---|---|---|---|---|
| (Adhikari et al., 2025b) | Feb-Nov 2023 | 2021-2025 | United States | integrated | farm | Farm | boot, sponge-stick swabs, flypapers, beetle traps, litter, soil, feed | environment | SE + PCR or WGS | 34 | 262 |
| | Feb-Nov 2023 | 2021-2025 | United States | integrated | processing plant | Slaughter facility | faces, boots | environment | SE + PCR or WGS | 4 | 12 |
| | Feb-Nov 2023 | 2021-2025 | United States | integrated | processing plant | Slaughter facility | post-pick whole carcass rinses | post-pick sample | SE + PCR or WGS | 39 | 64 |
| | Feb-Nov 2023 | 2021-2025 | United States | integrated | processing plant | Slaughter facility | post-chill whole carcass rinses | post-chill sample | SE + PCR or WGS | 28 | 97 |
| | Feb-Nov 2023 | 2021-2025 | United States | integrated | transportation (cage) | Transportation | faces | environment | SE + PCR or WGS | 20 | 38 |
| (M. V. Zbrun et al., 2015) | not mention | unclear | Argentina | integrated | farm | Farm | cloacal swabs | broiler | SE + PCR or WGS | 40 | 180 |
| | not mention | unclear | Argentina | integrated | retail market | Retail | carcasses | product | SE + PCR or WGS | 22 | 60 |
| | not mention | 2011-2015 | Argentina | integrated | slaughterhouse | Slaughter facility | carcasses | product unclear | SE + PCR or WGS | 43 | 60 |
| (Zaidi et al., 2012) | 2003-2006 | before 2010 | Mexico | non-integrated | retail | Retail | raw retail meat | product | DR | 759 | 1302 |
| | 2003-2006 | before 2010 | Mexico | non-integrated | slaughterhouse | Slaughter facility | intestines (cecum) | broiler | DR | 1017 | 1087 |
| (Adhikari et al., 2024b) | Dec 2021 – Nov 2022 | 2021-2025 | United States | integrated | farm | Farm | boots, flypapers, beetle traps, feed, soil sample, water puddles, miscellaneous | environment | SE + PCR or WGS | 3 | 119 |
| | Dec 2021 – Nov 2022 | 2021-2025 | United States | integrated | processing plant | Slaughter facility | post-pick carcass rinses | post-pick sample | SE + PCR or WGS | 9 | 10 |
| (Al Hakeem et al., 2025) | 2014 - 2017 | not applicable | United States | non-integrated | farm | Farm | faces, soil | environment | DR + PCR | 578 | 1200 |
| | 2014 - 2017 | not applicable | United States | non-integrated | processing plant | Slaughter facility | whole carcass rinse | product unclear | DR + PCR | 69 | 200 |
| | 2014 - 2017 | not applicable | United States | non-integrated | retail | Retail | whole carcass rinse | product | DR + PCR | 34 | 220 |
| (Iannetti et al., 2020) | not mention | unclean | Italy | integrated | farm | Farm | cloacal swabs | broiler | SE + PCR or WGS | 315 | 400 |
| | not mention | unclear | Italy | integrated | slaughterhouse | Slaughter facility | caeca | broiler | SE + PCR or WGS | 175 | 225 |
| | not mention | unclear | Italy | integrated | slaughterhouse | Slaughter facility | neck skin at post-pick stage | post-pick sample | SE + PCR or WGS | 68 | 221 |

| | | | | | | | | | | |
|---|---|---|---|---|---|---|---|---|---|---|
| | not mention | unclear | Italy | integrated | slaughterhouse | Slaughter facility | neck skin at pre-chill stage | pre-chill sample | SE + PCR or WGS | 260 | 444 |
| | not mention | unclear | Italy | integrated | slaughterhouse | Slaughter facility | neck skin at post-chill stage | post-chill sample | SE + PCR or WGS | 129 | 218 |
| (Van Asselt et al., 2008) | 2002-2005 | before 2010 | Netherlands | non-integrated | farm | Farm | feces | environment | DR | 1204 | 10266 |
| | 2002-2005 | before 2010 | Netherlands | non-integrated | processing plant | Slaughter facility | cecal samples | broiler | DR | 2486 | 10032 |
| | 2002-2005 | before 2010 | Netherlands | non-integrated | processing plant | Slaughter facility | breast skin | Product unclear | DR | 3434 | 15408 |
| (Carron et al., 2018) | Not report | unclear | Kenya | non-integrated | farm | Farm | faces, litter | environment | SE + PCR or WGS | 69 | 180 |
| | Not report | unclear | Kenya | non-integrated | retailer | Retail | raw chicken meat | product | SE + PCR or WGS | 68 | 181 |
| (Y. Tang et al., 2020) | not mention | 2016-2020 | China | integrated | farm | Farm | cloacal swabs | broiler | DR + PCR | 90 | 150 |
| | not mention | 2016-2020 | China | integrated | farm | Farm | sole, bedding, feed, water, floor, net, stool | environment | NR | 4 | 124 |
| | not mention | 2016-2020 | China | integrated | slaughterhouse | Slaughter facility | whole chicken carcasses at pre-pick stage | pre-pick sample | DR + PCR | 60 | 145 |
| | not mention | 2016-2020 | China | integrated | slaughterhouse | Slaughter facility | whole chicken carcasses after defeathering | post-pick sample | DR + PCR | 26 | 201 |
| | not mention | 2016-2020 | China | integrated | slaughterhouse | Slaughter facility | whole chicken carcasses after evisceration | pre-chill sample | DR + PCR | 102 | 191 |
| | not mention | 2016-2020 | China | integrated | slaughterhouse | Slaughter facility | whole chicken carcasses after cooling | post-chill sample | DR + PCR | 27 | 183 |
| | not mention | 2016-2020 | China | integrated | slaughterhouse | Slaughter facility | swab of machine surface | environment | NR | 24 | 88 |
| (Kojima et al., 2015) | Apr -Oct 2013 | 2011-2015 | Japan | non-integrated | slaughterhouse | Slaughter facility | meat, organs | product unclear | SE | 28 | 33 |
| | Apr -Oct 2013 | 2011-2015 | Japan | non-integrated | supermarket | Retail | meat, organs | product | SE | 22 | 75 |
| (Abdul Rahiman et al., 2023) | not mention | unclear | Malaysia | integrated | farm | Farm | boots, feeders, drinkers, faces | environment | SE | 313 | 460 |
| | not mention | unclear | Malaysia | integrated | processing plant | Slaughter facility | transport crates, stunning water | environment | SE | 134 | 170 |
| | not mention | unclear | Malaysia | integrated | retailer | Retail | wings, thighs, breasts | product | SE | 92 | 600 |
| (Damjanova et al., 2011) | Apr 2006 - Jan 2007 | before 2010 | Hungary | integrated | farm | Farm | faces | environment | SE + PCR or WGS | 57 | 84 |
| | Apr 2006 - Jan 2007 | before 2010 | Hungary | integrated | retail shop | Retail | raw chicken meat | product | SE + PCR or WGS | 45 | 55 |
| | Apr 2006 - Jan 2007 | before 2010 | Hungary | integrated | slaughterhouse | Slaughter facility | drag swab | environment | SE + PCR or WGS | 14 | 77 |

| | | | | | | | | | | | |
|---|---|---|---|---|---|---|---|---|---|---|---|
| | Apr 2006 - Jan 2007 | before 2010 | Hungary | integrated | slaughterhouse | Slaughter facility | post-chilled breast and neck skin | post-chill sample | SE + PCR or WGS | 63 | 70 |
| (M. V. Zbrun et al., 2017) | 2011-2012 | 2011-2015 | Argentina | integrated | farm | Farm | cloacal swabs | broiler | SE + PCR or WGS | 27 | 180 |
| | 2011-2012 | 2011-2015 | Argentina | integrated | farm | Farm | litter, feed and drinking water | environment | SE + PCR or WGS | 2 | 72 |
| | 2011-012 | 2011-2015 | Argentina | non-integrated | retail | Retail | carcasses | product | SE + PCR or WGS | 25 | 60 |
| | 2011-2012 | 2011-2015 | Argentina | integrated | slaughterhouse | Slaughter facility | carcasses after chilling | post-chill sample | SE + PCR or WGS | 17 | 60 |
| | 2011-2012 | 2011-2015 | Argentina | integrated | slaughterhouse | Slaughter facility | cecal | broiler | SE + PCR or WGS | 8 | 60 |
| | 2011-2012 | 2011-2015 | Argentina | integrated | slaughterhouse | Slaughter facility | liver | organ | SE + PCR or WGS | 5 | 60 |
| (Ramires et al., 2020) | Oct, Nov, and Dec 2014 | 2011-2015 | Brazil | non-integrated | farm | Farm | feces, drag swabs | environment | SE + PCR or WGS | 16 | 48 |
| | Oct, Nov, and Dec 2014 | 2011-2015 | Brazil | non-integrated | slaughterhouse | Slaughter facility | pre-pick samples (carcasses) | pre-pick sample | SE + PCR or WGS | 13 | 24 |
| | Oct, Nov, and Dec 2014 | 2011-2015 | Brazil | non-integrated | slaughterhouse | Slaughter facility | post-pick samples (carcasses) | post-pick sample | SE + PCR or WGS | 8 | 12 |
| | Oct, Nov, and Dec 2014 | 2011-2015 | Brazil | non-integrated | slaughterhouse | Slaughter facility | pre-chill samples (carcasses) | pre-chill sample | SE + PCR or WGS | 7 | 12 |
| | Oct, Nov, and Dec 2014 | 2011-2015 | Brazil | non-integrated | slaughterhouse | Slaughter facility | post-chill samples (carcasses) | post-chill sample | SE + PCR or WGS | 10 | 12 |
| | Oct, Nov, and Dec 2014 | 2011-2015 | Brazil | non-integrated | slaughterhouse | Slaughter facility | liver | organ | SE + PCR or WGS | 10 | 12 |
| | Oct, Nov, and Dec 2014 | 2011-2015 | Brazil | non-integrated | slaughterhouse | Slaughter facility | cecum | broiler | SE + PCR or WGS | 9 | 12 |
| | Oct, Nov, and Dec 20142014) | 2011-2015 | Brazil | non-integrated | slaughterhouse | Slaughter facility | scalding water, chiller water | environment | SE + PCR or WGS | 2 | 8 |
| (Kuria et al., 2017) | Oct 2012 - Sep 2013 | 2011-2015 | Kenya | non-integrated | farm | Farm | cloacal swabs | broiler | DR + PCR | 142 | 280 |
| | Oct 2012 - Sep 2013 | 2011-2015 | Kenya | non-integrated | live bird market | Live bird market | cloacal swabs | broiler | DR + PCR | 37 | 390 |
| | Oct 2012 - Sep 2013 | 2011-2015 | Kenya | non-integrated | market's slaughter facility | Slaughter facility | carcasses | product unclear | DR + PCR | 11 | 40 |
| (Meeyam et al., 2004) | may-Jul in 2000, 2001, and 2002 | before 2010 | Thailand | non-integrated | farm | Farm | cloacal swabs | broiler | DR + PCR | 156 | 425 |
| | may-Jul in 2000, 2001, and 2002 | before 2010 | Thailand | non-integrated | fresh chicken meat market | Retail | meat | product | DR + PCR | 27 | 72 |
| | may-Jul in 2000, 2001, and 2002 | before 2010 | Thailand | non-integrated | slaughterhouse | Slaughter facility | carcasses (after killing and defeathering of the chickens but before putting into the chilling tank) | pre-chill sample | DR + PCR | 64 | 352 |

| | | | | | | | | | | | |
|---|---|---|---|---|---|---|---|---|---|---|---|
| (M. Zbrun et al., 2013) | not mention | unclear | Argentina | integrated | farm | Farm | cloacal swabs | broiler | SE + PCR or WGS | 26 | 90 |
| | not mention | unclear | Argentina | integrated | retail market | Retail | carcasses | product | SE + PCR or WGS | 25 | 30 |
| | not mention | 2011-2015 | Argentina | integrated | slaughterhouse | Slaughter facility | carcasses after chilling | post-chill sample | SE + PCR or WGS | 10 | 30 |
| (Bort et al., 2022) | Jan 2016 -Dec 2017 | 2016-2020 | Spain | non-integrated | slaughterhouse | Slaughter facility | carcasses, neck skin after cooling | post-chill sample | SE | 98 | 430 |
| | Jan 2016 -Dec 2017 | 2016-2020 | Spain | non-integrated | supermarket | Retail | packaged broiler | product | SE | 23 | 48 |
| (Xin et al., 2025) | Mar 2021- Mar 2022 | 2021-2025 | Vietnam | non-integrated | farm | Farm | cecal contents | broiler | DR + PCR | 129 | 250 |
| | Mar 2021- Mar 2022 | 2021-2025 | Vietnam | non-integrated | live wholesale market | Live bird market | cecal contents | broiler | DR + PCR | 31 | 80 |
| | Mar 2021- Mar 2022 | 2021-2025 | Vietnam | non-integrated | slaughterhouse/sla ughter point | Slaughter facility | cecal contents | broiler | DR + PCR | 35 | 110 |
| (B. Tang et al., 2022) | Sep 2020 - Dec 2020 | 2016-2020 | China | non-integrated | farms | Farm | cloacal swabs | broiler | SE + MALDI-TOF | 29 | 30 |
| | Sep 2020 - Dec 2020 | 2016-2020 | China | non-integrated | slaughterhouse | Slaughter facility | cecum | broiler | SE + MALDI-TOF | 62 | 430 |
| | Sep 2020 - Dec 2020 | 2016-2020 | China | non-integrated | supermarket | Retail | meat | product | SE + MALDI-TOF | 20 | 257 |
| (Rossler et al., 2020) | May 2015 -Dec 2015 | 2011-2015 | Argentina | integrated | farm | Farm | cloacal swabs | broiler | SE + PCR or WGS | 106 | 318 |
| | May 2015 -Dec 2015 | 2011-2015 | Argentina | integrated | farm | Farm | boots, litter, feed, drinking water, files, darkling beetles | environment | SE + PCR or WGS | 12 | 281 |
| | May 2015 -Dec 2015 | 2011-2015 | Argentina | integrated | slaughterhouse | Slaughter facility | cecum | broiler | DR | 62 | 130 |
| | May 2015 -Dec 2015 | 2011-2015 | Argentina | integrated | slaughterhouse | Slaughter facility | post-chiller surfaces and washing of broiler carcasses | post-chill sample | DR | 23 | 145 |
| | May 2015 -Dec 2015 | 2011-2015 | Argentina | integrated | slaughterhouse | Slaughter facility | swab of surface, knives and worker hands, water | environment | SE | 29 | 156 |
| (Jozwiak et al., 2006) | Jun-July 2003 | before 2010 | Hungary | integrated | farm | Farm | cloacal swabs | broiler | SE | 1 | 10 |
| | Jun-July 2003 | Before 2010 | Hungary | integrated | farm | Farm | feed, litter, water, air, insects, and surface of the wall | environment | SE | 2 | 52 |
| | Jun-July 2003 | before 2010 | Hungary | integrated | slaughterhouse | Slaughter facility | cloaca sample | broiler | SE | 21 | 30 |
| | Jun-July 2003 | before 2010 | Hungary | integrated | slaughterhouse | Slaughter facility | surface of the carcass after plucking | post-pick sample | SE | 29 | 30 |

| | | | | | | | | | | | |
|---|---|---|---|---|---|---|---|---|---|---|---|
| | Jun-July 2003 | before 2010 | Hungary | integrated | slaughterhouse | Slaughter facility | surface of the carcass after evisceration | pre-chill sample | SE | 59 | 60 |
| | Jun-July 2003 | before 2010 | Hungary | integrated | slaughterhouse | Slaughter facility | surface of the carcass after chilling | post-chill sample | SE | 23 | 23 |
| | Jun-July 2003 | before 2010 | Hungary | integrated | slaughterhouse | Slaughter facility | personal hygiene, environmental hygiene | environment | SE | 32 | 33 |
| | Jun-July 2003 | Before 2010 | Hungary | integrated | transportation (cage) | Transportation | swab of animal transport boxes | environment | SE | 9 | 13 |
| (L. Ma et al., 2014) | Oct-Nov 2012 | 2011-2015 | China | integrated | farm | Farm | cecal samples | broiler | DR + PCR | 103 | 142 |
| | Oct-Nov 2012 | 2011-2015 | China | integrated | slaughterhouse | Slaughter facility | whole broiler carcasses at post-chill stage | post-chill sample | SE + PCR or WGS | 46 | 135 |
| | Oct-Nov 2012 | 2011-2015 | China | integrated | supermarket | Retail | whole broiler carcasses | product | SE + PCR or WGS | 40 | 128 |
| (Béjaoui et al., 2022) | 2017-2018 | 2016-2020 | Tunisia | non-integrated | retail store | Retail | carcasses | product | SE + PCR or WGS | 16 | 115 |
| | 2017-2018 | 2016-2020 | Tunisia | non-integrated | slaughterhouse | Slaughter facility | carcasses | product unclear | SE + PCR or WGS | 32 | 142 |

Abbreviations: SE: Selective enrichment; DP: Direct plating; PCR: Polymerase Chain Reaction; WGS: Whole Genome Sequencing

* It represents the number of samples for *Campylobacter* spp., including *C. coli*, *C. jejuni* and other *Campylobacter* spp.

**Supplementary Table S 7:** Summary of study characteristics of all included studies in the NTS group

| Study | Study year | Study year category | Country | PDN type | Sampling site | PDN component category | Sample source | Sample source | Detection method | Serovar identification method | No of positive samples | Sample size |
|---|---|---|---|---|---|---|---|---|---|---|---|---|
| (El Hage et al., 2022) | Oct 2014-Oct 2015 | 2011-2015 | Lebanon | non-integrated | slaughter plants | Slaughter facility | caeca | broiler | SE | agglutination | 55 | 115 |
| | Oct 2014-Oct 2015 | 2011-2015 | Lebanon | non-integrated | slaughter plants | Slaughter facility | post-chilling carcass | post-chill sample | SE | agglutination | 25 | 115 |
| | Oct 2014-Oct 2015 | 2011-2015 | Lebanon | non-integrated | commercial broiler farms | Farm | fecal samples | environment | SE | agglutination | 50 | 159 |
| | Oct 2014-Oct 2015 | 2011-2015 | Lebanon | non-integrated | retail shops | Retail | liver, whole chicken carcasses and cuts, and marinated chicken meat | product | SE | agglutination | 427 | 1907 |
| (Nelson et al., 2020) | May-Oct 2017 | 2016-2020 | Nepal | integrated | slaughterhouses | Slaughter facility | poultry tissue samples (muscle, skin, intestine, gizzard, heart, liver, spleen) from chicken carcass | product unclear | SE | NR | 126 | 159 |
| | May-Oct 2017 | 2016-2020 | Nepal | integrated | commercial poultry farms | Farm | environment samples (feces, litter, feed, water, vehicle swabs, inner walls) | environment | SE | NR | 31 | 62 |
| (Nhung et al., 2018) | Oct 2016-Mar 2017 | 2016-2020 | Vietnam | non-integrated | wet market | Retail | meat samples | product | SE + MALDI-TOF | MLST | 18 | 20 |
| | Oct 2016-Mar 2017 | 2016-2020 | Vietnam | non-integrated | supermarket | Retail | meat samples | product | SE + MALDI-TOF | MLST | 10 | 19 |
| (Wu et al., 2022) | Jul 2019-Sep 2020 | 2016-2020 | China | non-integrated | slaughterhouses | Slaughter facility | caecum and cloaca samples | broiler | SE + PCR | agglutination | 32 | 135 |
| | Jul 2019-Sep 2020 | 2016-2020 | China | non-integrated | slaughterhouses | Slaughter facility | environmental samples (swabs of desk, ground, instrument, and knife, shower water) | environment | SE + PCR | agglutination | 15 | 250 |
| | Jul 2019-Sep 2020 | 2016-2020 | China | non-integrated | farm | Farm | chicken cloaca | broiler | SE + PCR | agglutination | 7 | 188 |

| | | | | | | | | | | | | |
|---|---|---|---|---|---|---|---|---|---|---|---|---|
| | Jul 2019-Sep 2020 | 2016-2020 | China | non-integrated | farm | Farm | environmental samples (sewage, feed, chicken manure, fence, soli) | environment | SE + PCR | agglutination | 21 | 410 |
| | Jul 2019-Sep 2020 | 2016-2020 | China | non-integrated | retail markets | Retail | chicken meat | product | SE + PCR | agglutination | 7 | 86 |
| | Jul 2019-Sep 2020 | 2016-2020 | China | non-integrated | retail markets | Retail | environmental samples (swabs of chopping broad) | environment | SE + PCR | agglutination | 5 | 40 |
| (Adhikari et al., 2024b) | Dec 2021-Nov 2022 | 2021-2025 | United States | integrated | Processing | Slaughter facility | sponge-stick swabs, fly papers, miscellaneous | environment | SE | WGS | 2 | 40 |
| | Dec 2021-Nov 2022 | 2021-2025 | United States | integrated | farm | Farm | boot swabs, beetle traps, fly papers, litter grab, feed, soil samples, water samples, miscellaneous | environment | SE | WGS | 7 | 263 |
| | Dec 2021-Nov 2022 | 2021-2025 | United States | integrated | transport | Transportation | swab of the transport cages | environment | SE | WGS | 1 | 12 |
| (Pelyuntha et al., 2022) | Oct 2021-Mar 2022 | 2021-2025 | Thailand | non-integrated | slaughterhouses | Slaughter facility | swabs of food contact surfaces (working table, knife, cutting board, basket, and bucket), non-food contact surfaces (wall and floor), and equipment (conveyor) surfaces | environment | SE | agglutination | 18 | 102 |
| | Oct 2021-Mar 2022 | 2021-2025 | Thailand | non-integrated | commercial farms | Farm | animal feed, boot cover swabs, cooling pad water, drinking water, soil, and bedding material (rice husk) | environment | SE | agglutination | 9 | 102 |
| | Oct 2021-Mar 2022 | 2021-2025 | Thailand | non-integrated | free-range farm | Farm | animal feed, boot cover swabs, cooling | environment | SE | agglutination | 3 | 51 |

| | | | | | | | | | | | | |
|---|---|---|---|---|---|---|---|---|---|---|---|---|
| | | | | | | | pad water, drinking water, soil, and bedding material (rice husk) | | | | | |
| | Oct 2021-Mar 2022 | 2021-2025 | Thailand | non-integrated | chicken meat stalls | Retail | chicken meat | product | SE | agglutination | 21 | 153 |
| (Elgroud et al., 2009) | Mar 2005–Jun 2006 and Sep 2006–Mar 2007 | Before 2010 | Senegal | non-integrated | slaughterhouses | Slaughter facility | neck skin | product unclear | SE | agglutination | 12 | 150 |
| | Mar 2005–Jun 2006 and Sep 2006–Mar 2008 | Before 2010 | Senegal | non-integrated | slaughterhouses | Slaughter facility | environment samples (swabs of tables and knives) | environment | SE | agglutination | 13 | 540 |
| | Mar 2005–Jun 2006 and Sep 2006–Mar 2009 | Before 2010 | Senegal | non-integrated | broiler farms | Farm | faces and cloacal swabs | broiler | SE | agglutination | 7 | 450 |
| | Mar 2005–Jun 2006 and Sep 2006–Mar 2010 | Before 2010 | Senegal | non-integrated | broiler farms | Farm | faces and environment samples (feed, water, swabs of utensils and wall) | environment | SE | agglutination | 23 | 1350 |
| (Katsande et al., 2025) | Aug 2021–Dec 2023 | 2021-2025 | Zimbabwe | integrated | slaughterhouses | Slaughter facility | caeca contents | broiler | SE | WGS | 5 | 30 |
| | Aug 2021–Dec 2023 | 2021-2025 | Zimbabwe | integrated | slaughterhouses | Slaughter facility | swabs from defeathering and evisceration machines | environment | SE | WGS | 4 | 40 |
| | Aug 2021–Dec 2023 | 2021-2025 | Zimbabwe | integrated | slaughterhouses | Slaughter facility | post-chill carcass rinsates | post-chill sample | SE | WGS | 2 | 30 |
| | Aug 2021–Dec 2023 | 2021-2025 | Zimbabwe | integrated | farms | Farm | fresh faces | broiler | SE | WGS | 3 | 80 |
| | Aug 2021–Dec 2023 | 2021-2025 | Zimbabwe | integrated | farms | Farm | feed, wall dust, boot swabs | environment | SE | WGS | 11 | 208 |
| | Aug 2021–Dec 2023 | 2021-2025 | Zimbabwe | integrated | retail markets | Retail | chilled chicken cuts | product | SE | WGS | 4 | 20 |
| (Castellanos et al., 2018) | 2008- 2013 | Not applicable | Colombia | non-integrated | slaughterhouses | Slaughter facility | cecal content and carcass rinse (broiler and product) | product unclear | SE | WGS | 140 | 644 |

| | | | | | | | | | | | | |
|---|---|---|---|---|---|---|---|---|---|---|---|---|
| | 2008- 2013 | Not applicable | Colombia | non-integrated | farm | Farm | fecal material and fresh feces | broiler | SE | WGS | 28 | 74 |
| | 2008- 2013 | Not applicable | Colombia | non-integrated | retail | Retail | carcass rinse and chicken thighs meat | product | SE | WGS | 410 | 1554 |
| (Chen et al., 2019) | Mar- Oct 2015 | 2011-2015 | China | non-integrated | slaughterhouses | Slaughter facility | unpacked fresh meat | product unclear | SE + PCR | agglutination | 14 | 150 |
| | Mar- Oct 2015 | 2011-2015 | China | non-integrated | slaughterhouses | Slaughter facility | swabs of knives, floor, tables | environment | SE + PCR | agglutination | 15 | 90 |
| | Mar- Oct 2015 | 2011-2015 | China | non-integrated | supermarket | Retail | unpacked fresh meat | product | SE + PCR | agglutination | 9 | 92 |
| (Parry-Hanson | not mention | not mention | Ghana | non-integrated | supermarket | Retail | chicken carcasses | product | SE + PCR | NR | 5 | 12 |
| Kunadu et al., 2020) | not mention | not mention | Ghana | non-integrated | live bird markets | Live bird market | chicken carcasses | product | SE + PCR | NR | 28 | 36 |
| (Molina et al., 2024) | Aug-Dec 2019 | 2016-2020 | Costa Rica | integrated | slaughter facility | Slaughter facility | caeca samples | broiler | SE + Sero | MLST | 65 | 171 |
| | Aug-Dec 2019 | 2016-2020 | Costa Rica | integrated | retail (supermarkets, butchers and central markets) | Retail | chicken meat samples | product | SE + Sero | MLST | 38 | 65 |
| (Vinueza-Burgos et al., | Not mention | not mention | Ecuador | integrated | slaughterhouse | Slaughter facility | caecum | broiler | SE | ERIC-PCR | 2 | 5 |
| 2019) | Not mention | not mention | Ecuador | integrated | slaughterhouse | Slaughter facility | skin from pre-chilling carcass | prechill sample | SE | ERIC-PCR | 22 | 25 |
| | Not mention | not mention | Ecuador | integrated | slaughterhouse | Slaughter facility | skin from post-chilling carcass | post-chill sample | SE | ERIC-PCR | 14 | 25 |
| | Not mention | not mention | Ecuador | integrated | broiler farms | Farm | overshoes | environment | SE | ERIC-PCR | 27 | 51 |
| (Tőzsér et al., 2019a) | Feb-Apr 2017 | 2016-2020 | Hungary | non-integrated | slaughterhouse | Slaughter facility | body surface (after evisceration) | prechill sample | SE + PCR | NR | 9 | 10 |
| | Feb-Apr 2017 | 2016-2020 | Hungary | non-integrated | slaughterhouse | Slaughter facility | organ (offal) | organ | SE + PCR | NR | 5 | 5 |
| | Feb-Apr 2017 | 2016-2020 | Hungary | non-integrated | retail | Retail | wings, thighs, drumsticks, breasts, back | product | SE + PCR | NR | 30 | 30 |
| (Bellil et al., 2025) | May 2021-Jul 2022 | 2021-2025 | Algeria | non-integrated | slaughterhouses | Slaughter facility | intestines | broiler | SE + MALDI-TOF | WGS | 3 | 153 |
| | May 2021-Jul 2022 | 2021-2025 | Algeria | non-integrated | slaughterhouses | Slaughter facility | Internal organs | organ | SE + MALDI-TOF | WGS | 1 | 153 |
| | May 2021-Jul 2022 | 2021-2025 | Algeria | non-integrated | retail markets and buchers | Retail | raw poultry meat, chicken carcasses | product | SE + MALDI-TOF | WGS | 6 | 234 |

| | | | | | | | | | | | | |
|---|---|---|---|---|---|---|---|---|---|---|---|---|
| (Cui et al., 2016) | Aug-Sep, 2013 | 2011-2015 | China | integrated | abattoir (slaughterhouse) | Slaughter facility | Pre-chilling carcass | prechill sample | SE + PCR | agglutination | 28 | 203 |
| | Aug-Sep, 2013 | 2011-2015 | China | integrated | broiler farms | Farm | rectal swabs | broiler | SE + PCR | agglutination | 33 | 290 |
| | Aug-Sep, 2013 | 2011-2015 | China | integrated | retail markets | Retail | whole carcass | product | SE + PCR | agglutination | 29 | 126 |
| (Dione et al., 2009) | not mention | not mention | Senegal | non-integrated | broiler farms | Farm | fresh fecal samples | broiler | SE | agglutination | 20 | 57 |
| | not mention | not mention | Senegal | non-integrated | sale points | Retail | skin from chicken carcasses, meat from chicken carcasses | product | SE | agglutination | 39 | 114 |
| (Cui et al., 2019) | Aug-Sep, 2013 | 2011-2015 | China | integrated | abattoir(slaughterhouse) | Slaughter facility | post-chilling carcass | post-chill sample | DR + PCR | agglutination | 26 | 99 |
| | Aug-Sep, 2013 | 2011-2015 | China | integrated | abattoir(slaughterhouse) | Slaughter facility | butchered carcass (leg or breast, segmented chicken) | post-chill sample | DR + PCR | agglutination | 33 | 80 |
| | Aug-Sep, 2013 | 2011-2015 | China | integrated | market | Retail | carcasses | product | DR + PCR | agglutination | 16 | 127 |
| (Choi et al., 2014) | Nov 2011-Mar 2012 | 2011-2015 | Korea | integrated | slaughterhouses | Slaughter facility | post-chilling carcass | post-chill sample | SE + PCR | agglutination | 15 | 20 |
| | Nov 2011-Mar 2012 | 2011-2015 | Korea | integrated | slaughterhouses | Slaughter facility | drag device | environment | SE + PCR | agglutination | 31 | 52 |
| | Nov 2011-Mar 2012 | 2011-2015 | Korea | integrated | farms | Farm | drag and dust samples | environment | SE + PCR | agglutination | 52 | 324 |
| | Nov 2011-Mar 2012 | 2011-2015 | Korea | integrated | supermarket | Retail | chicken meat | product | SE + PCR | agglutination | 20 | 129 |
| | Nov 2011-Mar 2012 | 2011-2015 | Korea | integrated | broiler trucks | Transportation | drag samples | environment | SE + PCR | agglutination | 5 | 7 |
| (Ren et al., 2016) | Jul-Dec 2013 | 2011-2015 | China | integrated | slaughterhouse | Slaughter facility | whole chicken carcasses | product unclear | SE | agglutination | 22 | 35 |
| | Jul-Dec 2013 | 2011-2015 | China | integrated | farm | Farm | sterile cloacal swabs | broiler | SE | agglutination | 21 | 300 |
| | Jul-Dec 2013 | 2011-2015 | China | integrated | retail stores | Retail | whole chicken carcasses | product | SE | agglutination | 41 | 75 |
| (Xin et al., 2025) | Mar 2021-Mar 2022 | 2021-2025 | Vietnam | non-integrated | slaughterhouse | Slaughter facility | boot socks or overshoes | environment | SE + PCR | WGS | 3 | 6 |
| | Mar 2021-Mar 2022 | 2021-2025 | Vietnam | non-integrated | slaughter point | Slaughter facility | boot socks or overshoes | environment | SE + PCR | WGS | 8 | 19 |
| | Mar 2021-Mar 2022 | 2021-2025 | Vietnam | non-integrated | farm | Farm | boot socks or overshoes | environment | SE + PCR | WGS | 9 | 50 |
| | Mar 2021-Mar 2022 | 2021-2025 | Vietnam | non-integrated | retail | Retail | boot socks or overshoes | environment | SE + PCR | WGS | 6 | 16 |

| | | | | | | | | | | | | |
|---|---|---|---|---|---|---|---|---|---|---|---|---|
| | Mar 2021-Mar 2022 | 2021-2025 | Vietnam | non-integrated | wholesale | Live bird market | boot socks or overshoes | environment | SE + PCR | WGS | 6 | 11 |
| (S. Ma et al., 2017) | Mar 2015-Feb 2016 | 2011-2015 | China | integrated | abattoirs | Slaughter facility | cotton swabs | environment | SE | agglutination | 28 | 110 |
| | Mar 2015-Feb 2016 | 2011-2015 | China | integrated | farm | Farm | rectal swabs | broiler | SE | agglutination | 24 | 109 |
| | Mar 2015-Feb 2016 | 2011-2015 | China | integrated | market | Retail | meat samples | product | SE | agglutination | 36 | 116 |
| (Elsayed, 2019) | Summer 2017 | 2016-2020 | Egypt | integrated | slaughterhouse | Slaughter facility | environment samples (swabs from knives, table, and abattoir wall) | environment | SE | agglutination | 12 | 50 |
| | Summer 2017 | 2016-2020 | Egypt | integrated | slaughterhouse | Slaughter facility | pre-chilling carcass | prechill sample | SE | agglutination | 30 | 120 |
| | Summer 2017 | 2016-2020 | Egypt | integrated | slaughterhouse | Slaughter facility | chicken fillets | product unclear | SE | agglutination | 10 | 25 |
| | Summer 2017 | 2016-2020 | Egypt | integrated | slaughterhouse | Slaughter facility | livers | organ | SE | agglutination | 10 | 25 |
| | Summer 2017 | 2016-2020 | Egypt | integrated | farms | Farm | cloacal swabs | broiler | SE | agglutination | 55 | 145 |
| | Summer 2017 | 2016-2020 | Egypt | integrated | farms | Farm | feed, litter, water, farm walls, hand swabs from workers | environment | SE | agglutination | 35 | 75 |

Abbreviations: SE: Selective enrichment; DP: Direct plating; PCR: Polymerase Chain Reaction; WGS: Whole Genome Sequencing; ERIC-PCR: Enterobacterial Repetitive Intergenic Consensus PCR; MLST: Multilocus Sequence Typing; NR: not reported.

*The top three serovars are listed. The remaining were grouped under 'Others' to maintain clarity. 'Untyped' refers to isolates that were not typed or could not be identified in the original studies. NR: The number of serotypes is not reported.

**Supplementary Table S 8:** Summary of the random-effects subgroup analysis of *Campylobacter* spp. prevalence in farms and slaughter facilities

| Site | Subgroup | Studies | Prevalence, % (95% CI, %) | Heterogeneity | | |
|---|---|---|---|---|---|---|
| | | | | $\tau^2$ | $I^2$ | P value (Q test) * |
| Farm | Environment sample | 12 | 16.2 (6.9-33.5) | 2.70 | 99.4 | < 0.001 |
| | Broiler sample | 12 | 46.4 (30.4-63.2) | 1.33 | 98.2 | < 0.001 |
| Slaughter facility | Environment sample | 7 | 43.7 (18.5-72.7) | 2.48 | 96.4 | < 0.001 |
| | Broiler sample | 9 | 50.4 (26.8-73.8) | 2.35 | 99.2 | < 0.001 |
| | Organ sample | 2 | 39.1 (1.3-97.0) | 7.62 | 94.9 | < 0.001 |
| | Pre-pick sample | 2 | 44.4 (34.2-55.1) | 0.03 | 26.1 | 0.245 |
| | Post-pick sample | 6 | 61.5 (26.7-87.5) | 3.03 | 97.2 | < 0.001 |
| | Pre-chill sample | 5 | 62.4 (25.2-89.1) | 3.08 | 99.0 | < 0.001 |
| | Post-chill sample | 11 | 44.8 (25.3-66.0) | 1.98 | 97.6 | < 0.001 |
| | Other product sample | 6 | 42.9 (22.2-66.5) | 1.39 | 98.1 | < 0.001 |

- P value (Q test): the p-value of Cochran's Q test for between-study heterogeneity

**Supplementary Table S 9:** Summary of the random-effects subgroup analysis of NTS prevalence in farms and slaughter facilities

| Site | Subgroup | Studies | Prevalence, % (95% CI, %) | Heterogeneity | | |
|---|---|---|---|---|---|---|
| | | | | $\tau^2$ | $I^2$ | P value (Q test) * |
| farm | Environment sample | 10 | 13.4 (6.4-26.1) | 2 | 97 | < 0.001 |
| | Broiler sample | 10 | 13.8 (6.4-27.3) | 1.77 | 96.2 | < 0.001 |
| Slaughter facility | Environment sample | 11 | 17.5 (9.1-31.1) | 1.44 | 93.2 | < 0.001 |
| | Broiler sample | 6 | 22.3 (8.9-45.9) | 1.62 | 95.9 | < 0.001 |
| | Organ sample | 3 | 25.2 (5.4-95.4) | 12.21 | 93.8 | < 0.001 |
| | Pre-chill sample | 4 | 54.3 (13.5-90) | 3.92 | 97.4 | < 0.001 |
| | Post-chill sample | 6 | 35 (17.6-57.5) | 1.16 | 92.5 | < 0.001 |
| | Other product sample | 6 | 31.9 (12.1-61.5) | 2.26 | 97.8 | < 0.001 |

- P value (Q test): the p-value of Cochran's Q test for between-study heterogeneity

**Supplementary Table S 10:** Summary of univariable meta-regression analyses of study-level moderators of *Campylobacter* spp. prevalence at the farm, slaughter-facility, and retail levels

| Category | Covariate | Coefficient (95% CI) | $p$>\|z\|* | $R^2$(%)* | p>$\chi^2$* | $I^2$* |
|---|---|---|---|---|---|---|
| Farm | PDN type (reference: non-integrated PDN) | | | 2.9% | 0.12 | 99.3% |
| | • Integrated PDN | -1.08 (-2.4 to 0.3) | 0.12 | | | |
| | Country (reference: high income countries) | | | Nil | 0.59 | 99.3% |
| | • Upper-middle income countries | 0.44 (-1.06 to 1.94) | 0.56 | | | |
| | • Lower-middle income countries | 1.14 (-1.07 to 3.36) | 0.31 | | | |
| | Year of study conducted (reference: 2011-2015) | | | Nil | 0.5 | 99.9% |
| | • 2016-2020 | 1.11 (-1 to 3.21) | 0.3 | | | |
| | • 2021-2025 | -0.56 (-2.84 to 1.72) | 0.63 | | | |
| | • Before 2010 | -0.14 (-2.11 to 1.82) | 0.88 | | | |
| Slaughter facility | PDN type (reference: non-integrated PDN) | | | Nil | 0.71 | 99.3% |
| | • Integrated PDN | 0.16 (-0.71 to 1.03) | 0.71 | | | |
| | Country (reference: high income countries) | | | 4.8% | 0.02 | 99.2% |
| | • Upper-middle income countries | -0.9 (-1.76 to -0.05) | 0.04 | | | |
| | • Lower-middle income countries | -1.51 (-3.21 to 0.19) | 0.08 | | | |
| | Year of study conducted (reference: 2011-2015) | | | 12.6% | 0.06 | 99.3% |
| | • 2016-2020 | -0.82 (-1.89 to 0.26) | 0.14 | | | |
| | • 2021-2025 | 0.17 (-1.23 to 1.56) | 0.82 | | | |
| | • Before 2010 | 1.23 (0.16 to 2.29) | 0.02 | | | |
| Retail | PDN type (reference: non-integrated PDN) | | | 0.9% | 0.27 | 96.9% |
| | • Integrated PDN | 0.66 (-0.52 to 1.84) | 0.27 | | | |
| | Country (reference: high income countries) | | | Nil | 0.96 | 97.5% |
| | Upper-middle income countries | -0.19 (-1.79 to 1.41) | 0.96 | | | |
| | Lower-middle income countries | -0.05 (-1.80 to 1.71) | 0.81 | | | |
| | Year of study conducted (reference: 2011-2015) | | | 0.1% | 0.43 | 95.9% |
| | 2016-2020 | -0.16 (-1.87 to 1.55) | 0.85 | | | |
| | 2021-2025 | -0.73 (-2.70 to 1.25) | 0.47 | | | |
| | Before 2010 | 1.09 (-0.68 to 2.87) | 0.23 | | | |

- The p > |z| values are coefficient-level tests; for categorical covariates, they compare each non-reference category with the reference category and indicate whether it differs significantly from the reference category.
- The p >$\chi^2$ is from the overall Wald test and assesses whether the covariate as a whole is significantly associated with the true effect size (the estimation of *Campylobacter* spp. prevalence).
- $R^2$ represents the proportion of between-study variance ($\tau^2$) explained by the covariate.

- Residual $I^2$ represents the percentage of residual variation attributable to between-study heterogeneity after adjustment for the covariate.

**Supplementary Table S 11:** Summary of univariable meta-regression analyses of study-level moderators of NTS prevalence at the farm, slaughter-facility, and retail levels

| Category | Covariate | Coefficient (95% CI) | *p*>\|z\|* | $R^2$(%)* | p>$\chi^2$* | $I^2$* |
|---|---|---|---|---|---|---|
| Farm | PDN type (reference: non-integrated PDN) | | | Nil | 0.37 | 96.6% |
| | • Integrated PDN | 0.53 (-0.63 to 1.69) | 0.38 | | | |
| | Country (reference: high income countries) | | | Nil | 0.66 | 96.5% |
| | • Lower-middle income countries | 0.96 (-1.18 to 3.10) | 0.38 | | | |
| | • Upper-middle income countries | 0.66 (-1.51 to 2.84) | 0.55 | | | |
| | Year of study conducted (reference: 2011-2015) | | | 26.3% | 0.02 | 95.4% |
| | • 2016-2020 | 0.33 (-1.13 to 1.79) | 0.66 | | | |
| | • 2021-2025 | -1.05 (-2.47 to 0.38) | 0.15 | | | |
| | • Before 2010 | -1.21 (-2.91 to 0.48) | 0.16 | | | |
| Slaughter facility | PDN type (reference: non-integrated PDN) | | | 14.5% | 0.03 | 95.7% |
| | • Integrated PDN | 1 (0.08 to 1.92) | 0.03 | | | |
| | Country (reference: high income countries) | | | 4.5% | 0.11 | 96% |
| | • Lower-middle income countries | -1.36 (-2.83 to 0.11) | 0.07 | | | |
| | • Upper-middle income countries | -1.53 (-2.97 to -0.08) | 0.04 | | | |
| | Year of study conducted (reference: 2011-2015) | | | 31.5% | 0.001 | 94.2% |
| | • 2016-2020 | 0.28 (-0.79 to 1.35) | 0.61 | | | |
| | • 2021-2025 | -1.44 (-2.57 to -0.31) | 0.01 | | | |
| | • Before 2010 | -1.75 (-3.29 to -0.21) | 0.03 | | | |
| Retail | PDN type (reference: non-integrated PDN) | | | Nil | 0.93 | 98% |
| | • Integrated PDN | 0.06 (-1.19 to 1.3) | 0.93 | | | |
| | Country (reference: high income countries) | | | 20.4% | 0.02 | 95.6% |
| | Lower-middle income countries | -0.46 (-2.17 to 1.24) | 0.59 | | | |
| | Upper-middle income countries | -1.77 (-3.38 to -0.16) | 0.03 | | | |
| | Year of study conducted (reference: 2011-2015) | | | Nil% | 0.27 | 96.1% |
| | 2016-2020 | 1.26 (-0.23 to2.74) | 0.10 | | | |
| | 2021-2025 | -0.60 (-2.23 to 1.03) | 0.47 | | | |
| | Before 2010 | 0.24 (-2.43 to 2.91) | 0.86 | | | |

- The p > |z| values are coefficient-level tests; for categorical covariates, they compare each non-reference category with the reference category and indicate whether it differs significantly from the reference category.
- The p>$\chi^2$ is from the overall Wald test and assesses whether the covariate as a whole is significantly associated with the true effect size (the estimation of NTS prevalence).
- $R^2$ represents the proportion of between-study variance ($\tau^2$) explained by the covariate.

- Residual $I^2$ represents the percentage of residual variation attributable to between-study heterogeneity after adjustment for the covariate.

Supplementary Table S 12: Summary of all serovars and the corresponding isolates in farm, retail, and slaughter facilities

| Serovar | Farm | Slaughter facility | Retail | Total |
|---|---|---|---|---|
| Aarhus | - | 2 | 1 | 3 |
| Aequatoria | 1 | - | - | 1 |
| Agona | 7 | 16 | 19 | 42 |
| Albany | 4 | 7 | 4 | 15 |
| Anatum | - | 4 | 1 | 5 |
| Apeyeme | - | - | 1 | 1 |
| Bargny | 2 | 10 | - | 12 |
| Barranquilla | 4 | - | - | 4 |
| Blockley | 3 | 1 | 3 | 7 |
| Braenderup | 1 | 1 | 1 | 3 |
| Brancaster | 1 | 1 | - | 2 |
| Branderup | - | - | 2 | 2 |
| Bredeney | - | 3 | 1 | 4 |
| Calabar | - | - | 1 | 1 |
| Carnac | 4 | - | - | 4 |
| Cerro | - | 1 | - | 1 |
| Corvallis | - | 2 | 11 | 13 |
| Derby | 6 | 26 | 19 | 51 |
| Doncaster | - | - | 1 | 1 |
| Dublin | 2 | 1 | 1 | 4 |
| Emek | 4 | - | 3 | 7 |
| Enteritidis | 71 | 106 | 135 | 312 |
| Give | - | 5 | 1 | 6 |
| Glostrup | - | - | 1 | 1 |
| Gueuletapee | - | 7 | 9 | 16 |
| Hadar | 45 | 57 | 17 | 119 |
| Haifa | - | - | 1 | 1 |
| Heidelberg | 1 | - | 6 | 7 |
| I 4, [5],12:i:- | - | 1 | 1 | 2 |
| Indiana | 2 | 33 | 15 | 50 |
| Infantis | 36 | 70 | 215 | 321 |
| Istanboul | - | 2 | - | 2 |
| Jerusalem | - | 1 | 1 | 2 |
| Kedougou | - | - | 1 | 1 |
| Kentucky | 38 | 58 | 111 | 207 |
| Kiel | - | 1 | - | 1 |
| Labadi | - | 5 | - | 5 |
| Larochelle | - | 2 | - | 2 |
| Liverpool | 1 | 2 | - | 3 |
| Lomita | - | - | 1 | 1 |
| London | 3 | 4 | 2 | 9 |
| Luciana | 1 | - | - | 1 |
| Mbandaka | 11 | 5 | 4 | 20 |
| Meleagridis | 24 | 1 | 19 | 44 |
| Molade | 7 | 1 | - | 8 |
| Montevideo | - | 2 | 2 | 4 |
| Munster | - | - | 3 | 3 |
| Newport | - | 5 | 18 | 23 |
| Ohio | - | 1 | - | 1 |
| Papuana | 3 | - | - | 3 |
| Rideau | - | 2 | - | 2 |
| Rissen | 1 | - | 5 | 6 |
| Saintpaul | - | - | 4 | 4 |
| Schwarzengrund | - | 25 | 6 | 31 |
| Senftenberg | 8 | 4 | 2 | 14 |
| Singapore | - | 3 | - | 3 |
| Stanley | - | 1 | 2 | 3 |
| Stanleyville | - | 4 | - | 4 |
| Takoradi | 6 | - | - | 6 |
| Tamale | 3 | - | - | 3 |
| Thompson | 1 | - | 1 | 2 |
| Toulon | - | 1 | - | 1 |

| Serovar | Farm | Slaughter facility | Retail | Total |
|---|---|---|---|---|
| Typhimurium | 21 | 40 | 26 | 87 |
| Virchow | 8 | 9 | - | 17 |
| Virginia | - | - | 1 | 1 |
| Weltevreden | 22 | 15 | 14 | 51 |

**Supplementary Table S 13:** Summary of quality assessment for all included studies in the *Campylobacter* spp. group

| Study | Year | Sample strategy | Description of study subjects and setting | Coverage of sample type | Validity of micro-biological identification (*Campylobacter* spp.) | Reliability and Consistency of Measurement |
|---|---|---|---|---|---|---|
| (Abdul Rahiman et al., 2023) | 2022 | ☹ | ☺ | ☺ | ☹ | ☺ |
| (Adhikari et al., 2025a) | 2023 | ☺ | ☺ | ☺ | ☺ | ☺ |
| (Adhikari et al., 2024a) | 2021 | ☺ | ☺ | ☺ | ☺ | ☹ |
| (Béjaoui et al., 2022) | 2017-2018 | ☺ | ☺ | ☺ | ☺ | ☺ |
| (Bort et al., 2022) | 2016-2017 | ☹ | ☺ | ☺ | ☺ | ☺ |
| (Carron et al., 2018) | unknown | ☺ | ☺ | ☺ | ☺ | ☺ |
| (Damjanova et al., 2011) | 2006-2007 | ☺ | ☺ | ☺ | ☺ | ☹ |
| (Iannetti et al., 2020) | unknown | ☺ | ☺ | ☺ | ☺ | ☹ |
| (Jozwiak et al., 2006) | 2003-2004 | ☺ | ☺ | ☺ | ☹ | ☺ |
| (Kojima et al., 2015) | 2013 | ☹ | ☺ | ☺ | ☹ | ☹ |
| (Kuria et al., 2017) | 2012-2013 | ☹ | ☺ | ☺ | ☺ | ☺ |
| (L. Ma et al., 2014) | 2012 | ☹ | ☺ | ☺ | ☺ | ☺ |
| (Meeyam et al., 2004) | 2000-2003 | ☹ | ☹ | ☺ | ☺ | ☺ |
| (Ramires et al., 2020) | 2014 | ☹ | ☺ | ☺ | ☺ | ☺ |
| (Rossler et al., 2020) | 2015 | ☺ | ☺ | ☺ | ☺ | ☺ |
| (B. Tang et al., 2022) | 2020 | ☺ | ☹ | ☺ | ☺ | ☺ |
| (Y. Tang et al., 2020) | 2018 | ☺ | ☺ | ☺ | ☺ | ☺ |
| (Xin et al., 2025) | 2021-2022 | ☺ | ☺ | ☺ | ☺ | ☺ |
| (Zaidi et al., 2012) | 2003-2006 | ☹ | ☺ | ☺ | ☹ | ☺ |
| (M. V. Zbrun et al., 2015) | unknown | ☺ | ☹ | ☺ | ☺ | ☹ |
| (M. Zbrun et al., 2013) | unknown | ☺ | ☹ | ☺ | ☺ | ☺ |
| (M. V. Zbrun et al., 2017) | 2011-2012 | ☺ | ☺ | ☺ | ☺ | ☺ |
| (Al Hakeem et al., 2025) | 2014-2017 | ☹ | ☺ | ☺ | ☺ | ☺ |
| (Van Asselt et al., 2008) | 2002-2005 | ☺ | ☹ | ☺ | ☹ | ☹ |
| | | 15 | 19 | 24 | 20 | 18 |

**Supplementary Table S 14:** Summary of quality assessment for all included studies in the NTS group

| Study | Year | Sample strategy | Description of study subjects and setting | Coverage of sample type | Validity of microbiological identification (NTS) | Reliability and Consistency of Measurement |
|---|---|---|---|---|---|---|
| (Adel et al., 2021) | 2021 | ☺ | ☺ | ☺ | ☺ | ☺ |
| (Bellil et al., 2025) | 2021-2022 | ☹ | ☹ | ☺ | ☺ | ☺ |
| (Castellanos et al., 2018) | 2008-2013 | ☹ | ☹ | ☺ | ☹ | ☺ |
| (Chen et al., 2019) | 2015 | ☹ | ☺ | ☺ | ☺ | ☺ |
| (Choi et al., 2014) | 2011-2012 | ☹ | ☺ | ☺ | ☺ | ☺ |
| (Cui et al., 2016) | 2013 | ☹ | ☺ | ☺ | ☺ | ☺ |
| (Cui et al., 2019) | unknown | ☹ | ☹ | ☺ | ☺ | ☺ |
| (Dione et al., 2009) | unknown | ☺ | ☹ | ☺ | ☺ | ☺ |
| (Elgroud et al., 2009) | 2005-2007 | ☹ | ☺ | ☺ | ☺ | ☺ |
| (El Hage et al., 2022) | 2014-2016 | ☺ | ☺ | ☺ | ☺ | ☺ |
| (Elsayed, 2019) | 2017 | ☹ | ☺ | ☺ | ☺ | ☺ |
| (Katsande et al., 2025) | 2021-2023 | ☹ | ☺ | ☺ | ☺ | ☺ |
| (S. Ma et al., 2017) | 2015-2016 | ☹ | ☺ | ☺ | ☺ | ☹ |
| (Nhung et al., 2018) | 2016-2017 | ☹ | ☹ | ☺ | ☺ | ☺ |
| (Pelyuntha et al., 2022) | 2021-2022 | ☹ | ☺ | ☺ | ☺ | ☺ |
| (Ren et al., 2016) | 2013 | ☹ | ☹ | ☺ | ☺ | ☺ |
| (Vinueza-Burgos et al., 2019) | unknown | ☺ | ☺ | ☺ | ☺ | ☺ |
| (Wu et al., 2022) | 2019-2020 | ☹ | ☺ | ☺ | ☺ | ☺ |
| (Xin et al., 2025) | 2021-2022 | ☺ | ☺ | ☺ | ☺ | ☺ |
| (Parry-Hanson Kunadu et al., 2020) | unknown | ☺ | ☺ | ☹ | ☹ | ☺ |
| (Molina et al., 2024) | 2019 | ☹ | ☺ | ☺ | ☹ | ☺ |
| (Nelson et al., 2020) | 2017 | ☹ | ☺ | ☺ | ☹ | ☺ |
| (Tőzsér et al., 2019b) | 2017 | ☹ | ☺ | ☺ | ☹ | ☺ |
| | | 6 | 17 | 22 | 18 | 22 |

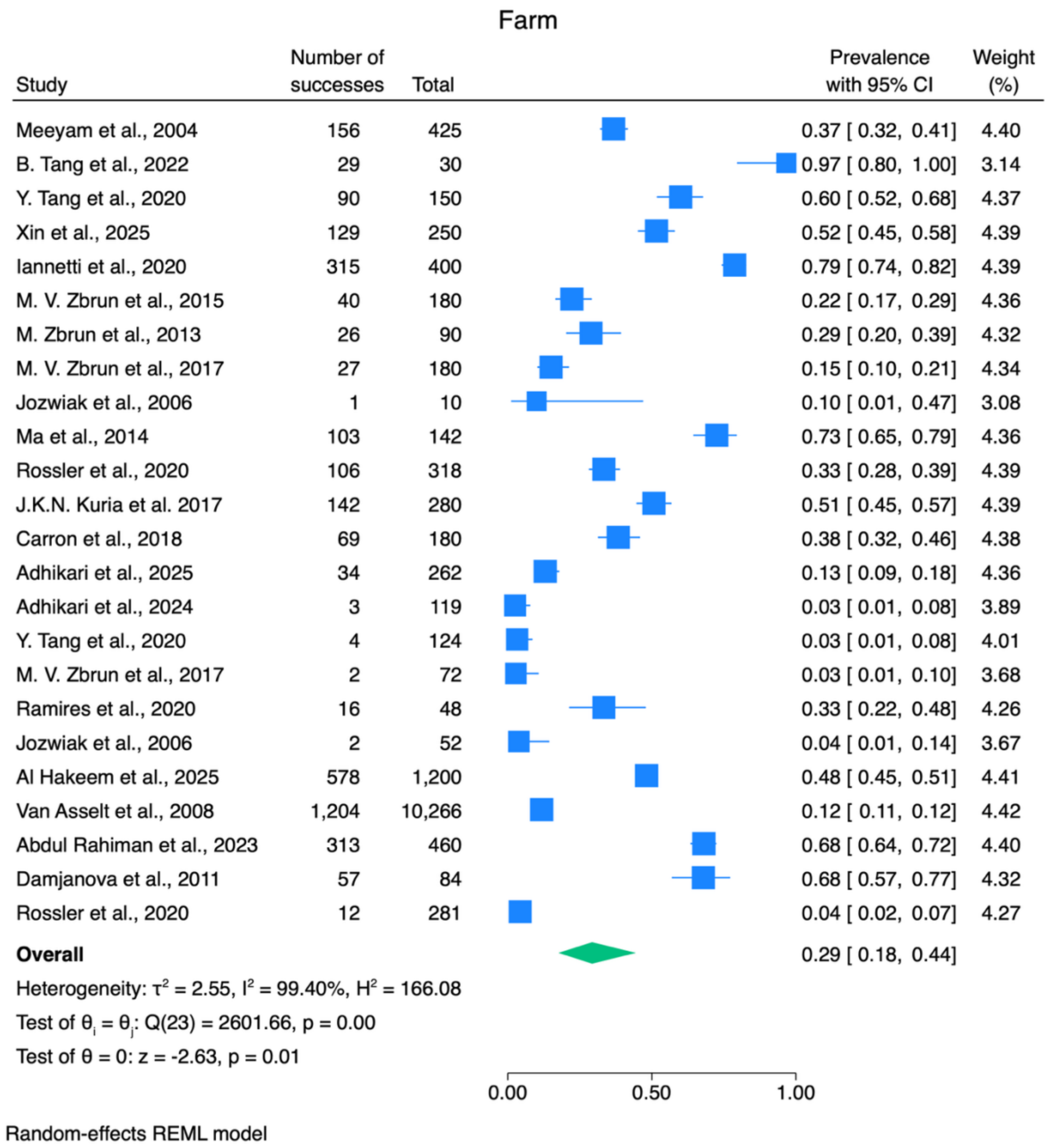

**Supplementary Figure S 1:** Mean prevalence of *Campylobacter* spp. in samples collected from farms

The number of successes and total represent the number of positive and total samples from each included study. $I^2$ represents the percentage of total variation due to between-study heterogeneity; $\tau^2$ represents the between-study variance; $Q$ and p-values are from Cochran's Q test for between-study heterogeneity.

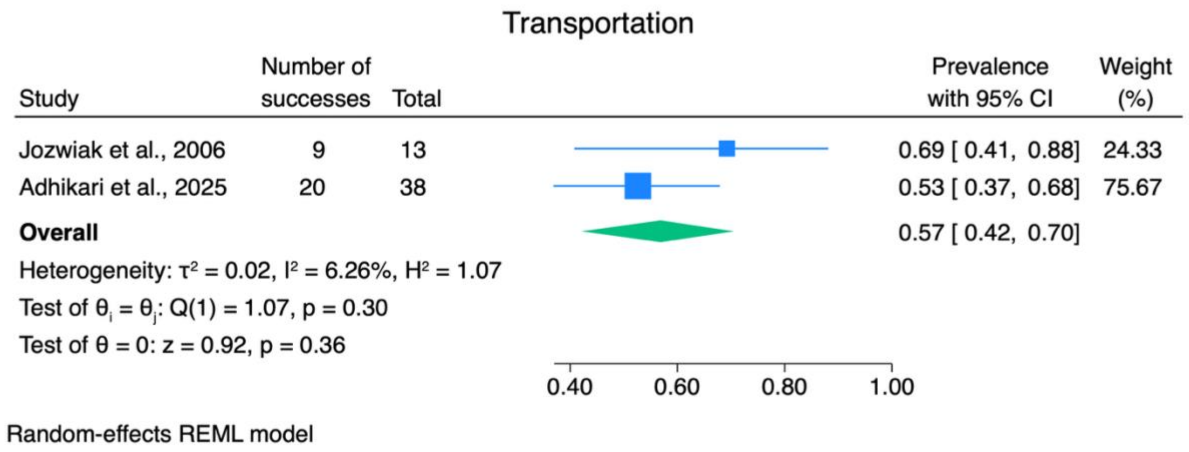

**Supplementary Figure S 2:** Mean prevalence of *Campylobacter* spp. in samples collected from transportation trucks

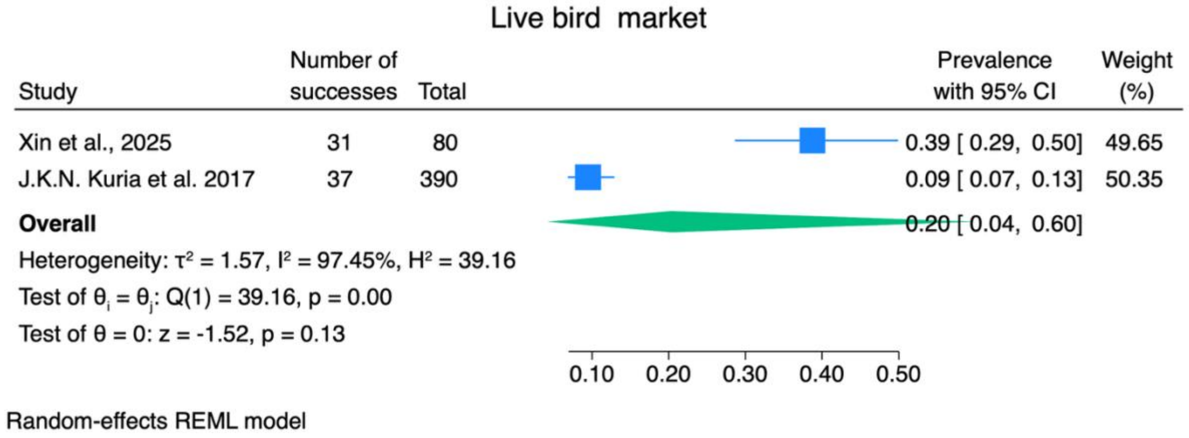

**Supplementary Figure S 3:** Mean prevalence of *Campylobacter* spp. in samples collected from live bird markets

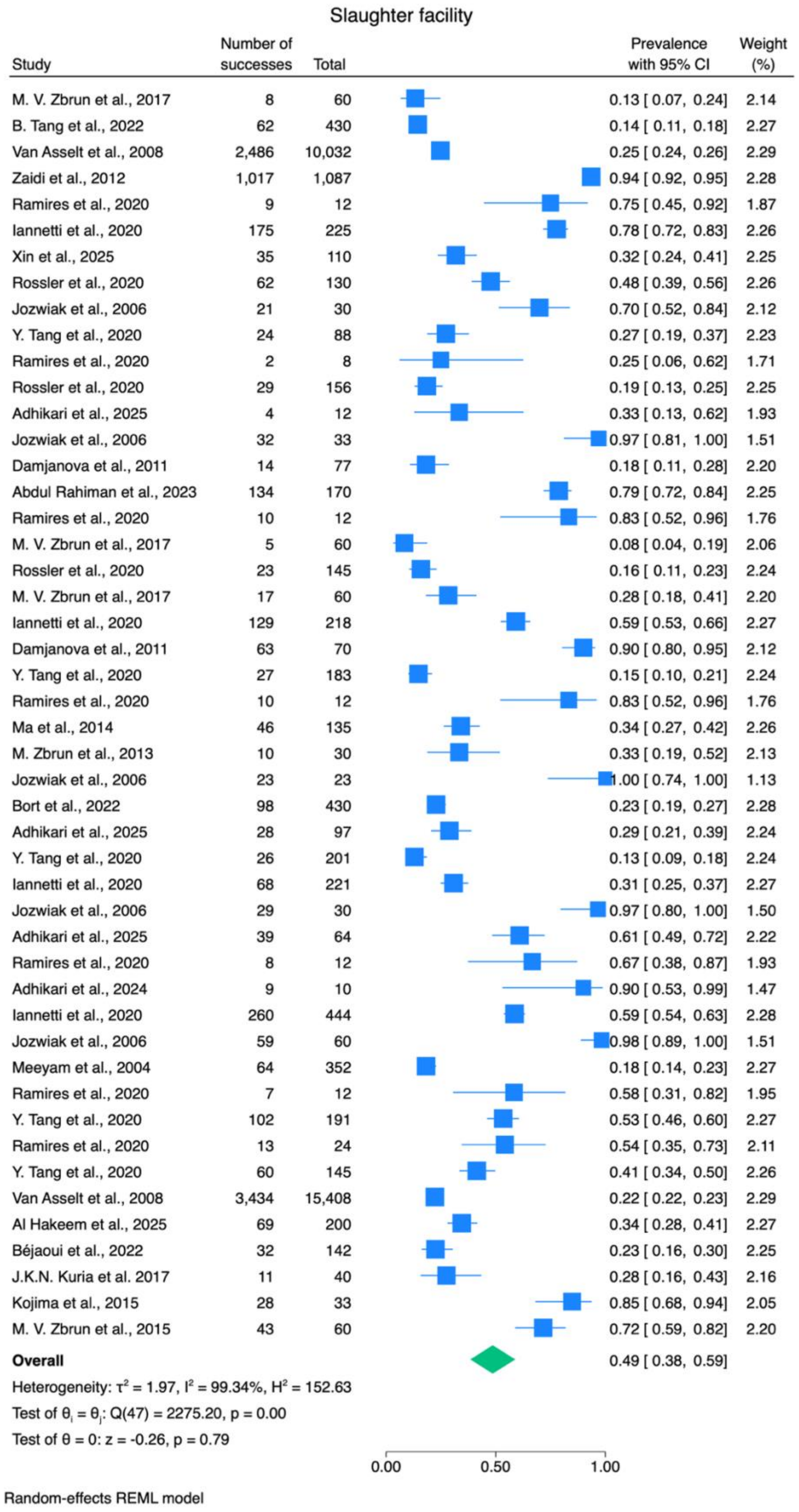

**Supplementary Figure S 4:** Mean prevalence of *Campylobacter* spp. in samples collected from slaughter facilities

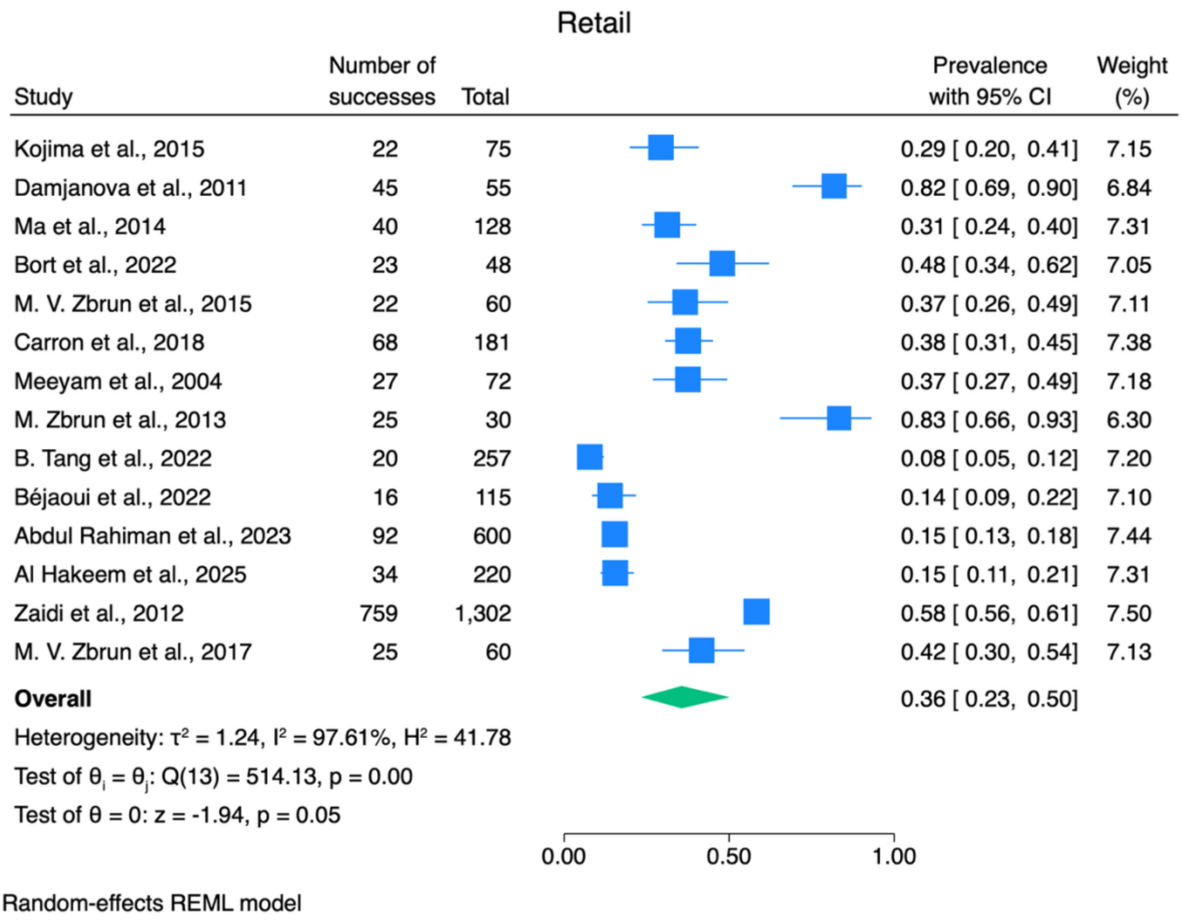

**Supplementary Figure S 5:** Mean prevalence of *Campylobacter* spp. in samples collected from retail

Farm

| Study | Number of successes | Total | Prevalence with 95% CI | Weight (%) |
|---|---|---|---|---|
| Adhikari et al., 2024 | 7 | 263 | 0.03 [ 0.01, 0.05] | 4.46 |
| Castellanos et al., 2018 | 28 | 74 | 0.38 [ 0.28, 0.49] | 4.67 |
| Choi et al., 2014 | 52 | 324 | 0.16 [ 0.12, 0.20] | 4.76 |
| Cui et al., 2016 | 33 | 290 | 0.11 [ 0.08, 0.16] | 4.73 |
| Dione et al., 2009 | 20 | 57 | 0.35 [ 0.24, 0.48] | 4.62 |
| Elgroud et al., 2009 | 7 | 450 | 0.02 [ 0.01, 0.03] | 4.46 |
| Elgroud et al., 2009 | 23 | 1,350 | 0.02 [ 0.01, 0.03] | 4.70 |
| El Hage et al., 2022 | 50 | 159 | 0.31 [ 0.25, 0.39] | 4.74 |
| Elsayed, 2019 | 55 | 145 | 0.38 [ 0.30, 0.46] | 4.74 |
| Elsayed, 2019 | 35 | 75 | 0.47 [ 0.36, 0.58] | 4.68 |
| Katsande et al., 2025 | 3 | 80 | 0.04 [ 0.01, 0.11] | 4.04 |
| Katsande et al., 2025 | 11 | 208 | 0.05 [ 0.03, 0.09] | 4.57 |
| Ma et al., 2017 | 24 | 109 | 0.22 [ 0.15, 0.31] | 4.68 |
| Pelyuntha et al., 2022 | 9 | 102 | 0.09 [ 0.05, 0.16] | 4.51 |
| Pelyuntha et al., 2022 | 3 | 51 | 0.06 [ 0.02, 0.17] | 4.03 |
| Ren et al., 2016 | 21 | 300 | 0.07 [ 0.05, 0.10] | 4.68 |
| Vinueza-Burgos et al., 2019 | 27 | 51 | 0.53 [ 0.39, 0.66] | 4.62 |
| Wu et al., 2022 | 7 | 188 | 0.04 [ 0.02, 0.08] | 4.45 |
| Wu et al., 2022 | 21 | 410 | 0.05 [ 0.03, 0.08] | 4.69 |
| Xin et al., 2025 | 9 | 50 | 0.18 [ 0.10, 0.31] | 4.48 |
| Parry-Hanson Kunadu et al., 2020 | 5 | 12 | 0.42 [ 0.18, 0.69] | 4.05 |
| Nelson et al., 2020 | 31 | 62 | 0.50 [ 0.38, 0.62] | 4.65 |
| **Overall** | | | 0.14 [ 0.08, 0.22] | |

Heterogeneity: $\tau^2 = 1.81$, $I^2 = 96.61\%$, $H^2 = 29.50$

Test of $\theta_i = \theta_j$: $Q(21) = 542.63$, $p = 0.00$

Test of $\theta = 0$: $z = -6.26$, $p = 0.00$

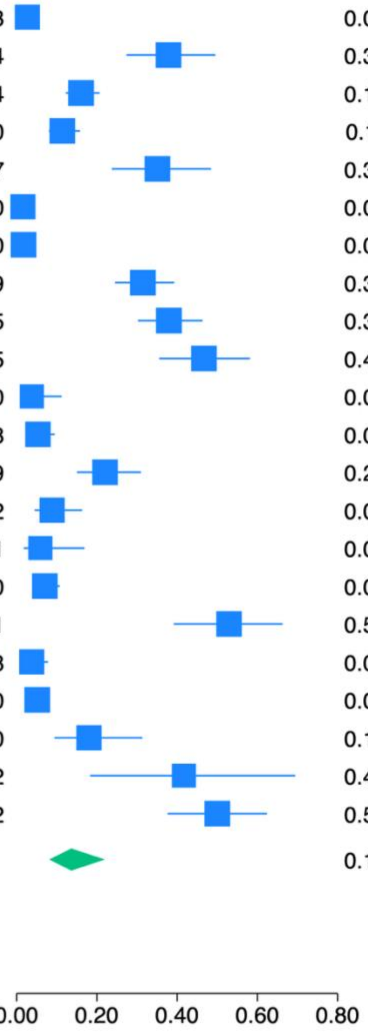

Random-effects REML model

**Supplementary Figure S 6:** Mean prevalence of NTS in samples collected from farms

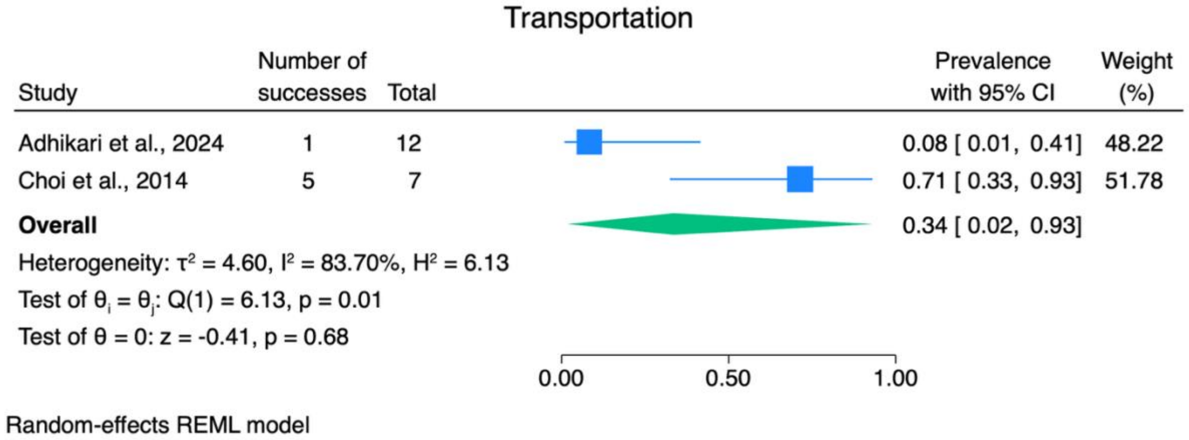

**Supplementary Figure S 7:** Mean prevalence of NTS in samples collected from transportation trucks

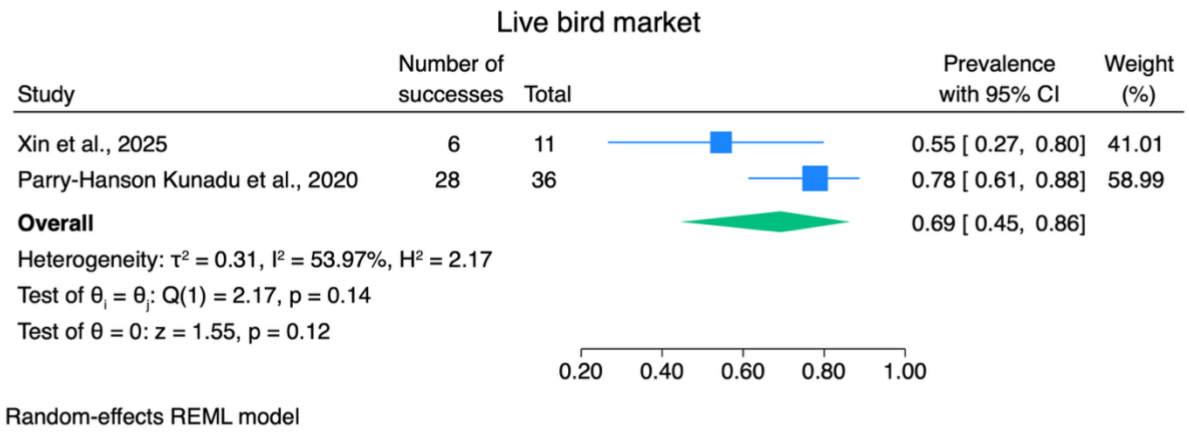

**Supplementary Figure S 8:** Mean prevalence of NTS in samples collected from live bird markets

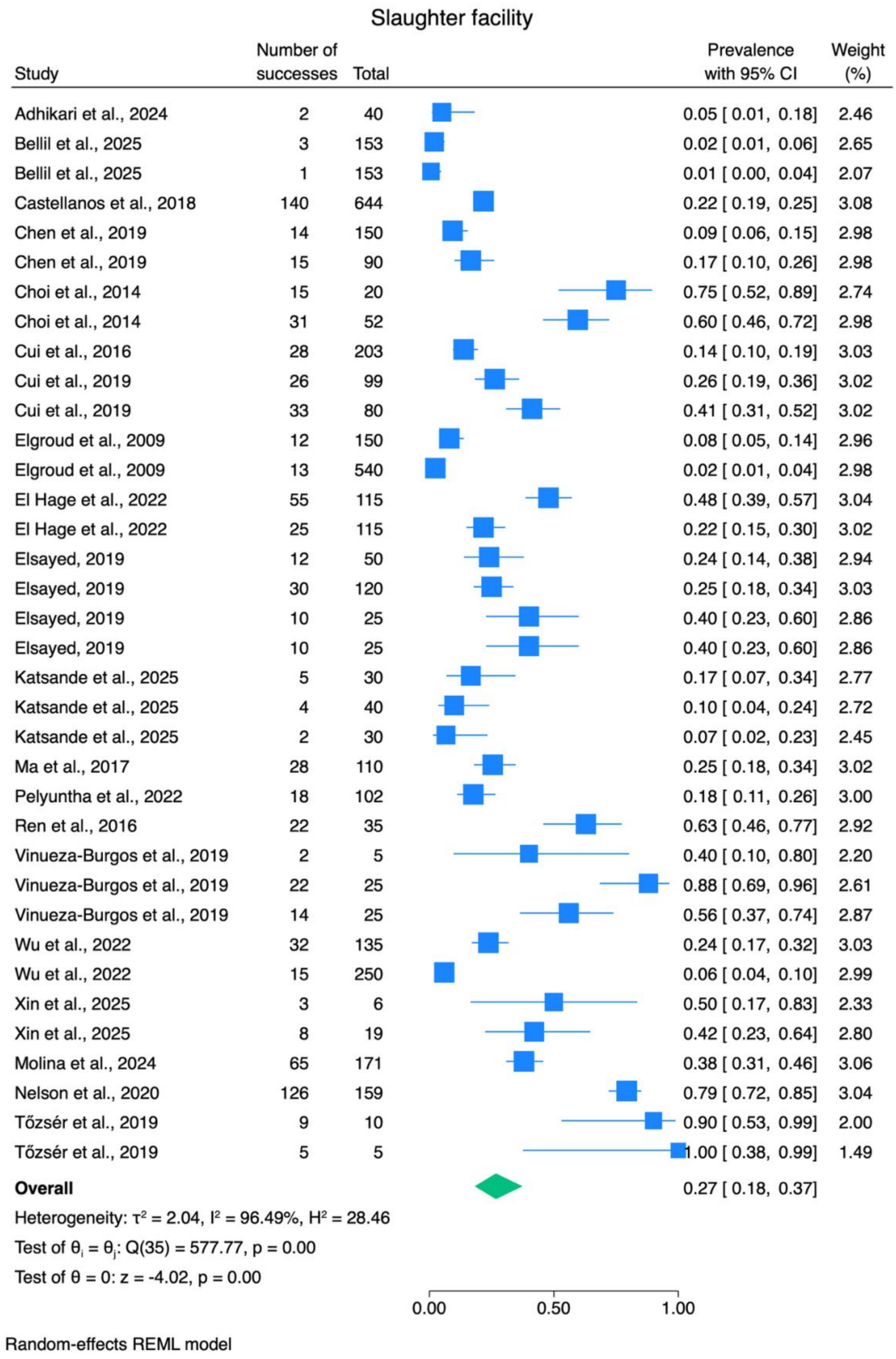

**Supplementary Figure S 9:** Mean prevalence of NTS in samples collected from slaughter facilities

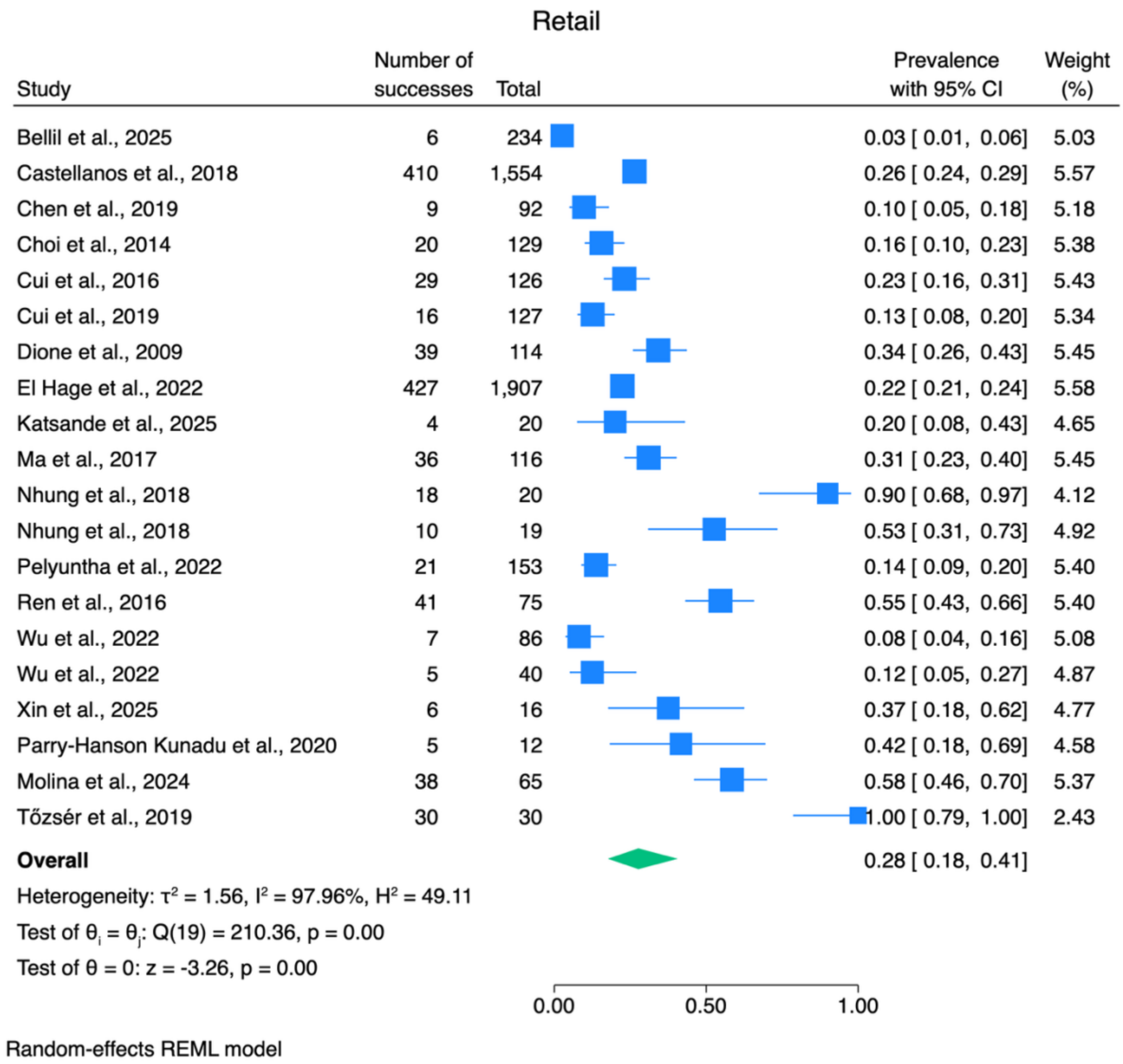

**Supplementary Figure S 10:** Mean prevalence of NTS in samples collected from retail

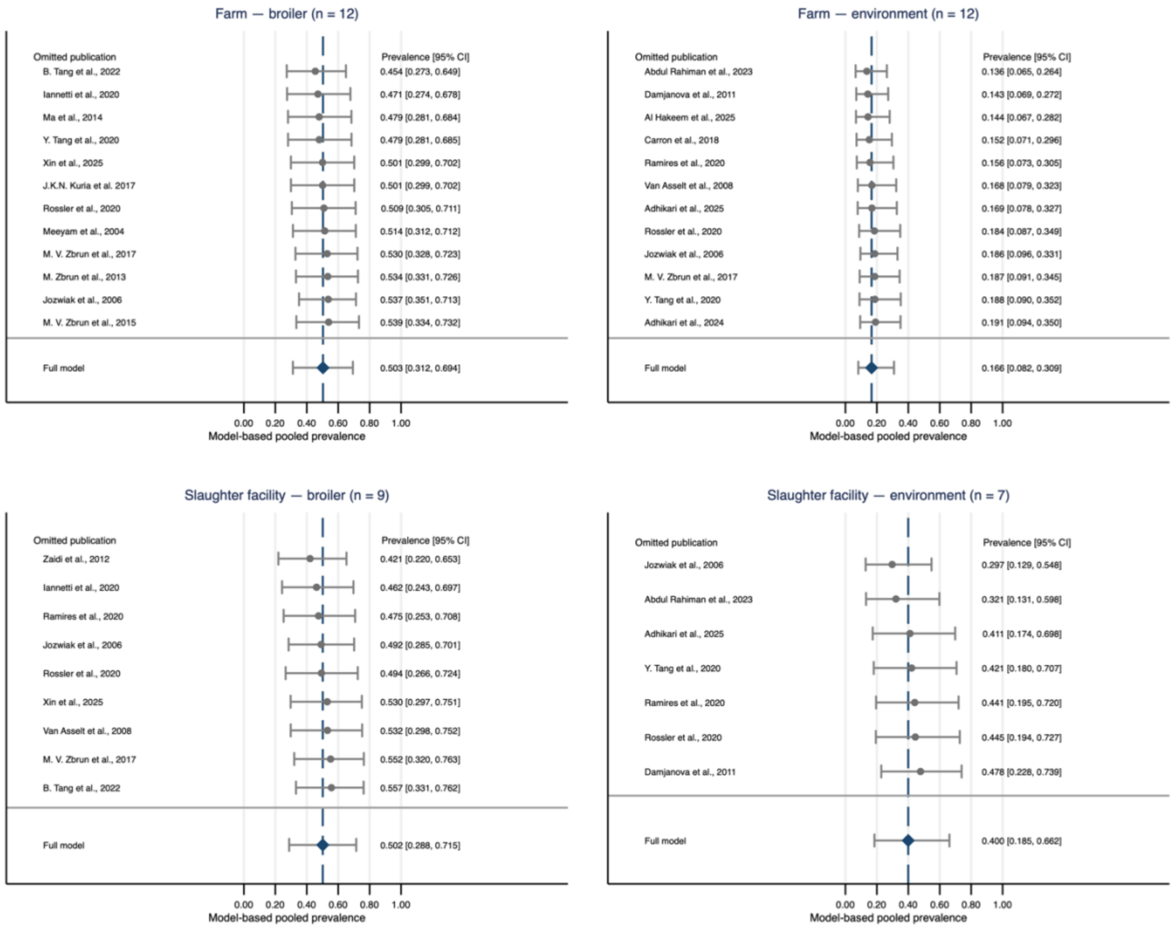

**Supplementary Figure S 11:** Leave-one-publication-out sensitivity analyses of pooled *Campylobacter* spp. prevalence in broiler and environmental samples from farms and slaughter facilities

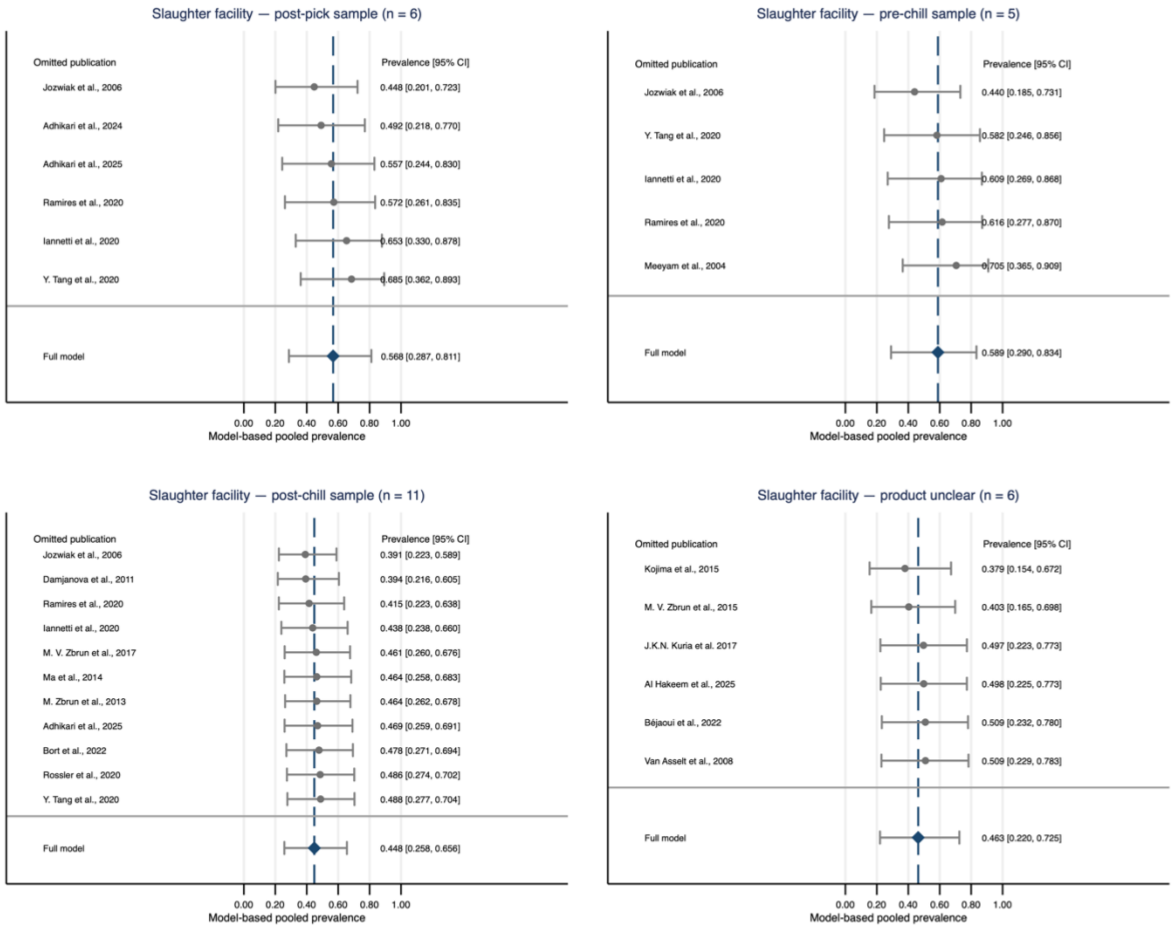

**Supplementary Figure S 12:** Leave-one-publication-out sensitivity analyses of pooled *Campylobacter* spp. prevalence in post-pick, pre-chill, post-chill, and other product samples without stage description, collected from slaughter facilities

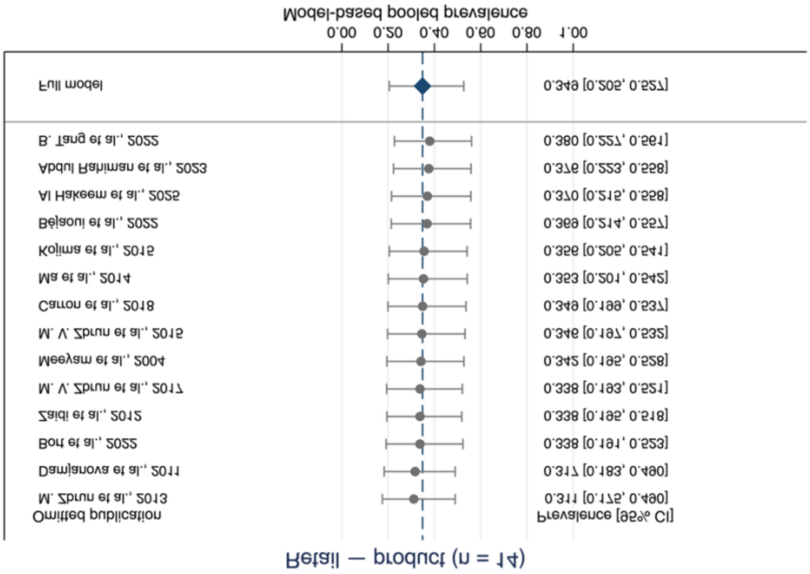

**Supplementary Figure S 13:** Leave-one-publication-out sensitivity analyses of pooled *Campylobacter* spp. prevalence in product samples collected from retail

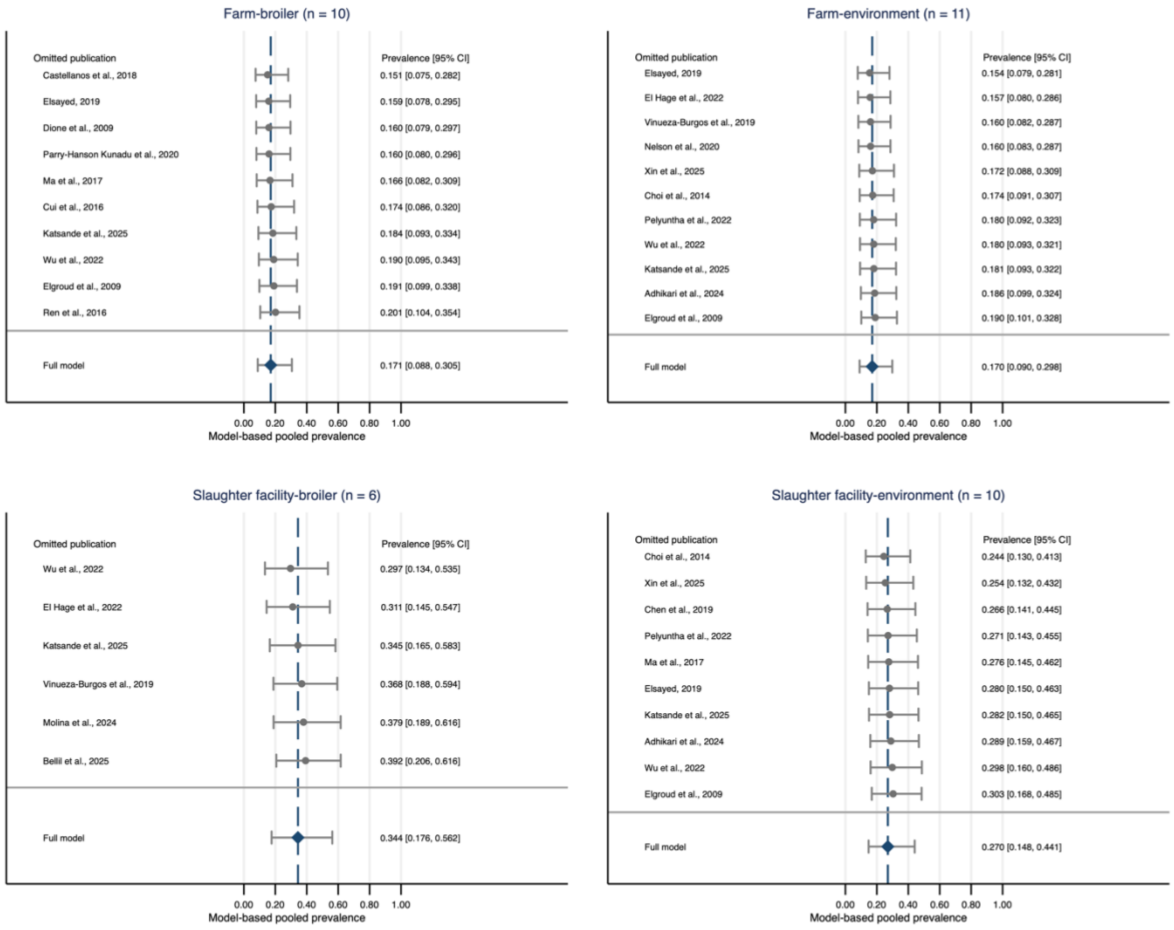

**Supplementary Figure S 14:** Leave-one-publication-out sensitivity analyses of pooled NTS prevalence in broiler and environmental samples from farms and slaughter facilities

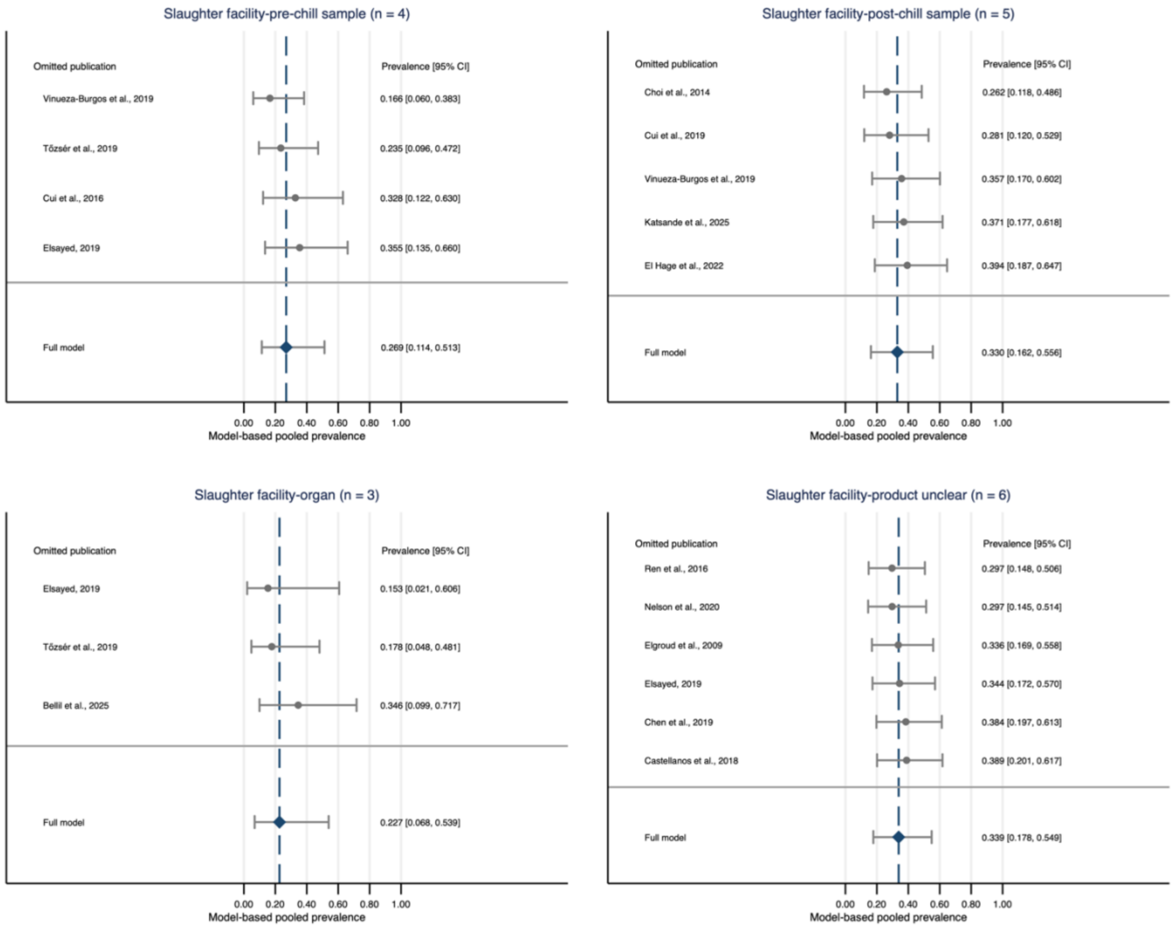

**Supplementary Figure S 15:** Leave-one-publication-out sensitivity analyses of pooled NTS prevalence in pre-chill, post-chill, organ, and other product samples without stage description, collected from slaughter facilities

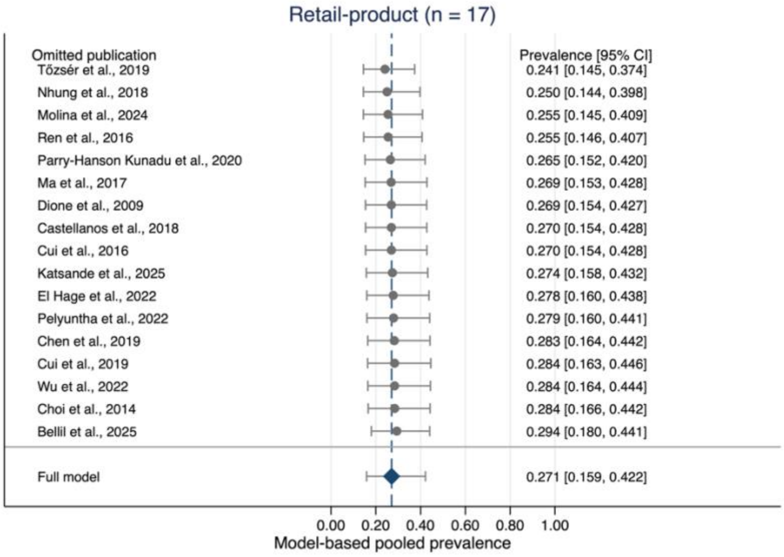

**Supplementary Figure S 16:** Leave-one-publication-out sensitivity analyses of pooled NTS prevalence in product samples collected from retail
